# GRAS-1 is a conserved novel regulator of early meiotic chromosome dynamics in *C. elegans*

**DOI:** 10.1101/2022.09.06.506874

**Authors:** Marina Martinez-Garcia, Pedro Robles Naharro, Marnie W. Skinner, Kerstin A. Baran, Saravanapriah Nadarajan, Nara Shin, Carlos G. Silva-García, Takamune T. Saito, Sara Beese-Sims, Ana Castaner, Sarai Pacheco, Enrique Martinez-Perez, Philip W. Jordan, Monica P. Colaiácovo

**Affiliations:** Department of Genetics, Blavatnik Institute, Harvard Medical School, Boston, MA, USA; Biochemistry and Molecular Biology Department, Johns Hopkins University, Bloomberg School of Public Health, Baltimore, MD, USA; Department of Biochemistry and Molecular Biology, Uniformed Services University of the Health Sciences, Bethesda, MD, USA; Department of Molecular Metabolism, Harvard T. H. Chan School of Public Health, Harvard University, Boston, MA, USA; MRC London Institute of Medical Sciences, London, UK

**Keywords:** GRAS-1, CYTIP, Tamalin, GRASP, meiosis, germline, chromosome dynamics, homologous pairing, synapsis, licensing, DNA double-strand break repair

## Abstract

Chromosome movements and licensing of synapsis must be tightly regulated during early meiosis to ensure accurate chromosome segregation and avoid aneuploidy, although how these steps are coordinated is not fully understood. Here we show that GRAS-1, the worm homolog of mammalian GRASP/Tamalin and CYTIP, coordinates early meiotic events with cytoskeletal forces outside the nucleus. GRAS-1 localizes close to the nuclear envelope (NE) in early prophase I and interacts with NE and cytoskeleton proteins. Delayed homologous chromosome pairing, synaptonemal complex (SC) assembly, and DNA double-strand break repair progression are partially rescued by the expression of human CYTIP in *gras-1* mutants, supporting functional conservation. However, *Tamalin, Cytip* double knockout mice do not exhibit obvious fertility or meiotic defects, suggesting evolutionary differences between mammals. *gras-1* mutants show accelerated chromosome movement during early prophase I, implicating GRAS-1 in regulating chromosome dynamics. GRAS-1-mediated regulation of chromosome movement is DHC-1-dependent, placing it acting within the LINC-controlled pathway, and depends on GRAS-1 phosphorylation at a C-terminal S/T cluster. We propose that GRAS-1 serves as a scaffold for a multi-protein complex coordinating the early steps of homology search and licensing of SC assembly by regulating the pace of chromosome movement in early prophase I.

## INTRODUCTION

Meiosis is a specialized cell division process in which diploid germ cells give rise to haploid gametes (i.e., eggs and sperm) accomplished by following a single round of DNA replication with two consecutive rounds of chromosome segregation. To segregate properly, homologous chromosomes must undergo a series of steps that are unique to the first meiotic division and are conserved from yeast to mammals, including: (1) pairing, (2) assembly of the “zipper-like” synaptonemal complex (SC) between paired homologs, and (3) formation of programmed meiotic DNA double-strand breaks (DSBs) resulting in crossover recombination, leading to genetic diversity and physical attachments between homologs (Láscarez-Lagunas et al. 2020). Errors in any of these steps can result in impaired chromosome segregation and aneuploidy, which is associated with 20% of birth defects (e.g., Down Syndrome), 35% of clinically recognized miscarriages, infertility, and tumorigenesis (Webster and Schuh 2017).

During pairing, homologous chromosomes must physically align along their lengths; this is achieved by pronounced chromosome movements inside the meiotic nucleus driven by cytoskeletal forces. In mammals and worms, this is achieved through the LINC (linker of nucleoskeleton and cytoskeleton) protein complex, which transmits forces to the nucleus/NE via cytoskeletal microtubules and dynein (Link and Jantsch 2019, Zetka et al. 2020). In *C. elegans*, the meiotic LINC complex is formed by the KASH-domain protein ZYG-12 at the outer nuclear membrane and the SUN-domain protein SUN-1 at the inner nuclear membrane (Cohen-Fix and Askjaer 2017). SUN-1 interacts via yet unidentified factor(s) with one end of each chromosome carrying specific repetitive sequences (pairing centers, PCs) which are bound by PC end Zinc-finger-proteins. PC proteins facilitate chromosome movement until homologs begin pairing and assembling the SC (Hillers et al. 2017). The SC is a tripartite structure composed of proteins assembled along chromosome axes (lateral elements) and proteins that bridge each pair of axes (central region components) (Lake and Hawley 2021). Studies in budding yeast, plants, flies, worms, and mammals, have shown that the SC is critical for stabilizing homologous chromosome pairing, the progression of meiotic recombination, crossover formation, and achieving accurate meiotic chromosome segregation (Zickler and Kleckner 2015). Work in *C. elegans* has identified proteins involved in regulating pairing and SC formation (Nadarajan et al. 2017; Alleva et al. 2017; Link et al. 2018; Bowman et al. 2019; Castellano-Pozo et al. 2020), but how NE-associated proteins regulate chromosome dynamics during early prophase I is incompletely understood. Here we show that *C. elegans* GRAS-1, which is homologous to mammalian GRASP/Tamalin and CYTIP, localizes to the NE and is required for the regulation of chromosome movement. GRAS-1 limits the speed of dynein-microtubule driving forces and contributes to the licensing of SC assembly, ensuring adequate timing of key meiotic processes such as homologous chromosome pairing, SC assembly, and DSB repair progression. While mice *Tamalin, Cytip* double knockout (DKO) mutants did not display obvious SC and recombination defects, human CYTIP partially rescued a *gras-1* mutation, supporting functional conservation and suggesting evolutionary differences between the mammalian proteins. We propose a model by which GRAS-1 links NE-cytoskeleton-SC assembly and coordinates early meiotic events by acting as a brake during early meiotic prophase I chromosome movements.

## RESULTS

### GRAS-1 localization is meiosis-specific and contacts NE components

A yeast-two hybrid screen for candidates interacting with worm SC proteins identified GRAS-1 (Smolikov et al. 2009). *gras-1* (ORF F30F8.3) encodes for a 245 amino acid protein containing PDZ (PSD-95/SAP90, Discs-large, and ZO-1) and coiled-coil (CC) domains (Fig. 1A). GRAS-1 shares a high degree of conservation with both human CYTIP (Cytohesin-interacting protein, 63% homology) and GRASP/Tamalin (General receptor for phosphoinositides 1-associated scaffold protein, 61% homology), due to a gene duplication event in chordates (Fig. 1A, B, TreeFam). No orthologs were found in fungi or plants. GRASP has been implicated as a scaffold for multi-protein complexes involved in processes such as epithelial cell migration and membrane trafficking (Kitano et al. 2002; Attar and Santy 2013). CYTIP plays roles in cell adhesion and the immune system (Heufler et al. 2008). In mice and humans, both GRASP and CYTIP are expressed in testes and ovaries (Fig. S1A) (Nevrivy et al. 2000; Uhlén et al. 2015; Human Protein Atlas). In worms, *gras-1* exhibits germline-enriched expression that is restricted to meiosis by the RNA-binding protein PUF-8 (Fig. S1B) (Kohara 2001; Reinke 2004; Ortiz et al. 2014; Tzur et al. 2018; Mainpal et al. 2011). However, the meiotic functions for GRAS-1 and its homologs remained unknown.

**Fig. 1.**
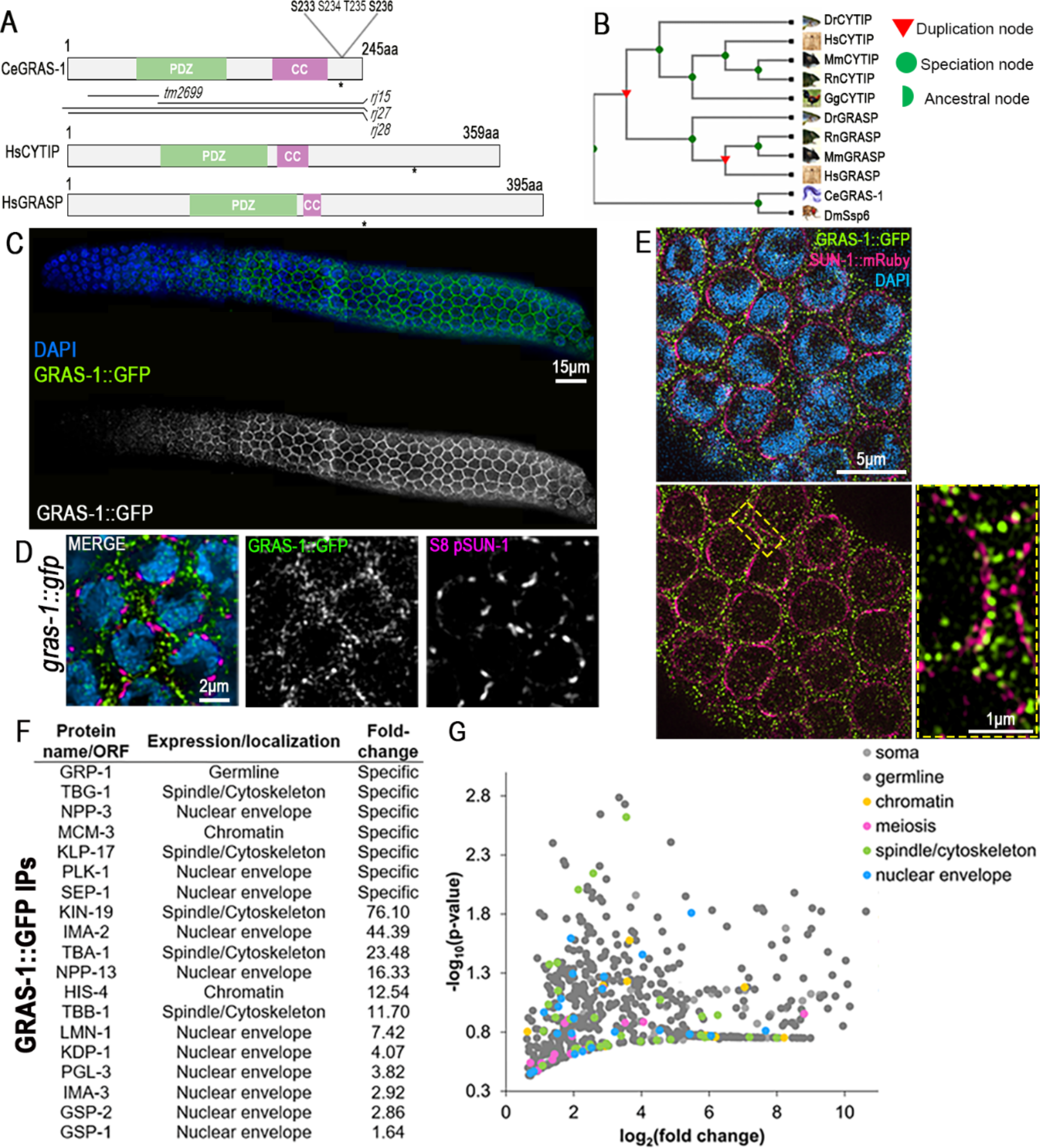
GRAS-1 localization is meiosis-specific and contacts NE components. **(A)** Protein conservation between *C. elegans* (Ce) GRAS-1 and *Homo sapiens* (Hs) CYTIP and GRASP. PDZ: PSD-95/SAP90, Discs-large and ZO-1, CC: coiled-coil, asterisk indicates position of predicted phosphorylation sites. **(B)** Evolutionary tree (TreeFam) of GRAS-1 orthologs in *Drosophila melanogaster*, *Mus musculus* (Mm), *Rattus norvegicus* (Rn), *Danio rerio* (Dr), *Gallus* (Gg). **(C)** GRAS-1::GFP localization in hermaphrodite gonads co-stained with anti-GFP (green) and DAPI (blue). **(D)** Higher magnification images of leptotene/zygotene stage nuclei co-stained with anti-GFP for GRAS-1::GFP (green), anti-S8 pSUN-1 (magenta) and DAPI (blue). **(E)** Super-resolution microscopy image of *gras-1::gfp* leptotene/zygotene nuclei co-stained for GRAS-1::GFP (green), SUN-1::mRuby (magenta) and DAPI. Dashed rectangle indicates region of the nuclear margins shown at higher magnification. **(F)** GRAS-1 interacting proteins. Immunoprecipitation from GRAS-1::GFP whole worm extracts was analyzed by mass spectrometry analysis. Their localization and enrichment in the MS samples compared to controls are shown. **(G)** Volcano plot depicting all MS analysis hits above a 1.5 fold-change in GRAS-1::GFP samples compared to controls (x axis), their statistical significance (y axis) and colored by their described expression/localization in *C. elegans*.

Different databases place GRAS-1 and its mammalian homologs at the plasma membrane, cytosol, membrane systems and the perinuclear region (WolFSORT, UniProt, Human Protein Atlas). The expression of a functional GRAS-1::GFP transgene revealed a meiosis-specific localization of GRAS-1 in both hermaphrodite and male germlines (Fig. 1C, S1C). GRAS-1::GFP signal was detected both at germ cell membranes, as confirmed by SYN-4 and Phalloidin staining (Fig. S1D), and near the nuclear envelope in early prophase I, as determined by co-immunolocalization with phosphorylated nuclear envelope protein SUN-1 (Fig. 1D; S8-pSUN-1). 73% of the nuclei in the leptotene/zygotene region had at least one S8-pSUN-1 aggregate contacting a GRAS-1::GFP signal (n=89) and 44% of the S8 pSUN-1 signals were in contact with GRAS-1::GFP (n=215, 13 gonads). Super-resolution microscopy analysis of a worm line expressing both SUN-1::mCherry and GRAS-1::GFP further supports GRAS-1 localization close to the NE (Fig. 1E). Moreover, GRAS-1::GFP localization appears to be largely independent of meiotic DSB production and SC formation (Fig. S1E). Using a transgenic line expressing GRAS-1-GFP for pull-downs and mass spectrometry analysis, we found proteins previously shown to be expressed in the germline (Fig. 1F). GRP-1 appeared as the most enriched protein in all 4 replicates and specific to the GRAS-1::GFP pull-downs. GRP-1 is the worm ortholog of human Cytohesin 1 protein, the main structural and functional partner of CYTIP (Heufler et al. 2008; Teuliere et al. 2014), supporting conservation between the proteins. Many of the proteins identified included NE-associated proteins, such as tubulins, PLK-1, importins, the KASH protein KDP-1, and cytoskeleton or spindle structural and motor components. Based on their GO terms or WormBase-described functions and/or localization, germline hits were classified into the following categories: nuclear envelope, spindle/cytoskeleton, meiosis, chromatin, or general germline-expressed proteins. The majority of proteins (667 out of 774, excluding GRAS-1) had a greater than 1.5 fold-change suggesting GRAS-1::GFP interactors are localized to/interact with the NE or cytoskeleton (Fig. 1G).

### GRAS-1 is required for normal meiotic progression and accurate chromosome segregation

To assess the roles of *gras-1* in the germline, we analyzed the fertility of various *gras-1* alleles including an out-of-frame deletion between the first and second exons (*tm2699*), a partial deletion and frameshift from amino acid 89 (*rj15)*, and whole-gene deletions (*rj27* and *rj28*) (Fig. 1A). While all mutants had normal brood sizes, most exhibited a mild but significant increase in the number of eggs laid that failed to hatch (embryonic lethality), elevated levels of male progeny (indicating meiotic chromosome nondisjunction), and increased larval lethality (Fig. S1F, 2A). To assess the effects of complete absence of GRAS-1 protein, all subsequent analyses were performed in *gras-1(rj28)* mutants.

Analysis of meiotic progression revealed an extension in the number of rows of nuclei exhibiting phosphorylated SUN-1 (S8 pSUN-1) signal in *gras-1* null mutants compared to wild type (22.8±0.6 and 19.5±0.5, respectively; p=0.0001, Mann Whitney U-test, Fig. 2B). This was accompanied by an increase in the number of rows of nuclei with chromosomes exhibiting the characteristic configuration of leptotene/zygotene stage nuclei in *C. elegans* (14±0.3 and 10.9±0.1, respectively; p<0.0001) (Fig. 2B). This alteration in meiotic progression is further supported by a delay in Polo-like kinase PLK-2 translocating from the nuclear periphery to synapsed chromosomes by the end of early pachytene (19±0.5 rows of nuclei in wild type and 21.9±1 in *gras-1*, p=0.0351) (Fig. S2A). These data suggest that GRAS-1 is required for normal meiotic progression and accurate chromosome segregation.

**Fig. 2.**
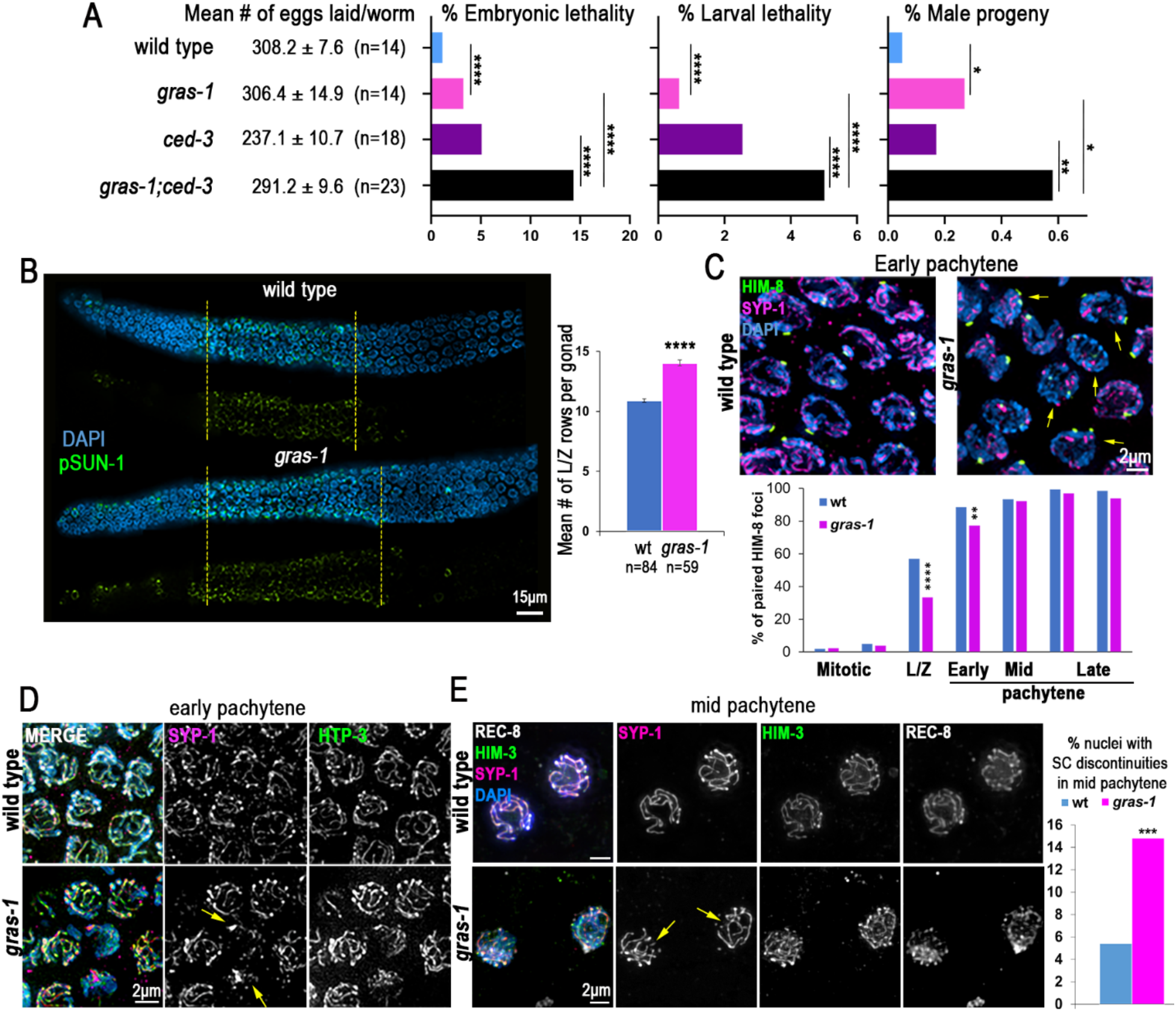
GRAS-1 is required for normal meiotic progression, chromosome pairing, and synapsis. **(A)** Mean number of eggs laid (brood size) ± SEM, the percentage of embryonic lethality, larval lethality, and male progeny are shown for the indicated genotypes. *p<0.05, **p=0.0037, ****p<0.0001 by Fisher’s exact test. n= number of P0 worms analyzed from three independent biological replicates. **(B)** Whole mounted gonads of wild type and *gras-1* worms co-stained with anti-S8 pSUN-1 (green) and DAPI (blue). Both merged and S8 pSUN-1 signal only are shown with yellow dotted lines delimiting the region in which complete rows of nuclei presented S8 pSUN-1 signal; n= 30 gonads each. Graph on the right shows the mean number of nuclei in leptotene/zygotene (L/Z) stage per gonad in wild type and *gras-1* worms. ****p<0.0001 by the Mann-Whitney U-test, n= number of worms analyzed from at least 2 independent biological replicates. **(C)** Top, high-resolution images of early pachytene nuclei co-stained with anti-HIM-8 (green), anti-SYP-1 (magenta) and DAPI (blue) from wild type and *gras-1* worms. Yellow arrows indicate nuclei with unpaired HIM-8 signal. Bottom, percentage of nuclei with paired HIM-8 signals (≤0.75μm apart) at different germline stages. ****p<0.0001, **p=0.008 by the Fisher’s Exact Test; n=6 gonads each and a minimum of 131 nuclei per zone. **(D)** High-resolution images of wild type and *gras-1* early pachytene nuclei (n= 80 and 50, respectively) from whole mounted gonads co-stained with anti-SYP-1 (magenta), anti-HTP-3 (green) and DAPI (blue). Yellow arrows indicate nuclei with SYP-1 aggregates. **(E)** Left, high-resolution images of lightly squashed gonads of wild type and *gras-1* mid pachytene nuclei co-stained with anti-SYP-1 (magenta), anti-HIM-3 (green), anti-REC-8 (white) and DAPI (blue). Right, percentage of mid-pachytene nuclei with SC discontinuities in wild type and *gras-1* gonads. ***p=0.0032, Fisher’s Exact test, n=253 and 217, respectively, from two biological replicates.

### GRAS-1 is necessary for timely homologous chromosome pairing and synapsis in an α-importin-independent manner

Because delays in meiotic progression during early prophase I can arise from problems in homolog pairing (Smolikov et al. 2007a; Sato et al. 2009; Alleva et al. 2017), we measured X chromosome pairing throughout meiosis by immunostaining for the X chromosome-specific PC protein HIM-8 (Phillips et al. 2005). During leptotene/zygotene and early pachytene stages, we observed higher levels of nuclei with two unpaired HIM-8 foci in *gras-1* mutants compared to wild type (p<0.0001 in leptotene/zygotene stage and p=0.008 in early pachytene, Fisher’s Exact Test) (Fig. 2C).

Early pachytene nuclei with unpaired HIM-8 foci in *gras-1* mutants also showed a discontinuous SC or aggregates of the SC central region protein SYP-1, in contrast to the continuous SC tracks detected in wild type (a mean of 2.28±0.24 compared to 0.4±0.08 nuclei with SYP-1 aggregates in *gras-1* and wild type, respectively; p<0.0001, Mann Whitney U-test) (Fig. 2D). Discontinuities of the central region of the SC, but not of axial element proteins such as HTP-3, were also detected along chromosomes in mid-pachytene nuclei of whole mounted germlines from *gras-1* mutants compared to wild type (16.9% and 5.1%, respectively; p=0.002, Fisher’s Exact test) (Fig. S2B) and further confirmed on squash preparations (14.8% and 5.4%, respectively; p=0.0032) (Fig. 2E).

The α-importin nuclear transport IMA-2 protein and the Akirin protein AKIR-1 have been proposed to act through parallel pathways to ensure normal chromosome synapsis by promoting import and chromosomal loading of cohesin complex proteins. For instance, *akir-1;ima-2* double mutants exhibit an increased number of nuclei with SC aggregates and discontinuities due to the abnormal loading of axis and cohesin proteins (Bowman et al. 2019). However, axial element proteins HTP-3 and HIM-3 and the meiosis-specific cohesin REC-8 were correctly loaded on the chromosomes in *gras-1* mutants, suggesting that SC complex defects may be caused by other mechanisms (Fig. 2D, E). Moreover, REC-8 localization was not altered in *gras-1* and *ima-1* or *ima-2*, double and triple mutants (Fig. S2C). Interestingly, we detected interaction of GRAS-1 with multiple SC central region proteins, including SYP-3 by western blot analysis of GRAS-1::GFP pull downs (Fig. S3A), and SYP-1, SYP-2, and SYP-3 by yeast two-hybrid analysis (Fig. S3B). Taken together, these studies support a role for GRAS-1 in promoting timely homologous chromosome pairing and SC assembly in an α-importin-independent manner during early prophase I.

### Early prophase I chromosome movement is limited by GRAS-1 in a dynein-dependent manner

The delay in homologous pairing and SC assembly observed in *gras-1* mutants is similar to that detected in mutants where chromosome movement is impaired (Wynne et al. 2012; Sato et al. 2009; Woglar et al. 2013). Therefore, we assessed chromosome movement by live imaging analysis of SUN-1::mRuby;GFP::H2B aggregates (marking chromosome ends) during meiosis in wild type and *gras-1* young adult worms. Surprisingly, SUN-1 aggregates moved at a greater speed and traveled higher distances in *gras-1* mutants compared to wild type (84.47±1.05 nm/s, average total distance traveled in 60s of 5.03±0.1 μm, and 50.55±0.82 nm/s, average total distance traveled in 60s of 2.99±0.07μm, respectively, p<0.0001 Student’s t-test) (Fig. 3A). Moreover, we did not observe increases in the area of the SUN-1 aggregates that could suggest more power to move chromosomes (0.184±0.006 μm in wild type and 0.179±0.006 μm in *gras-1*, p=0.5834).

**Fig. 3.**
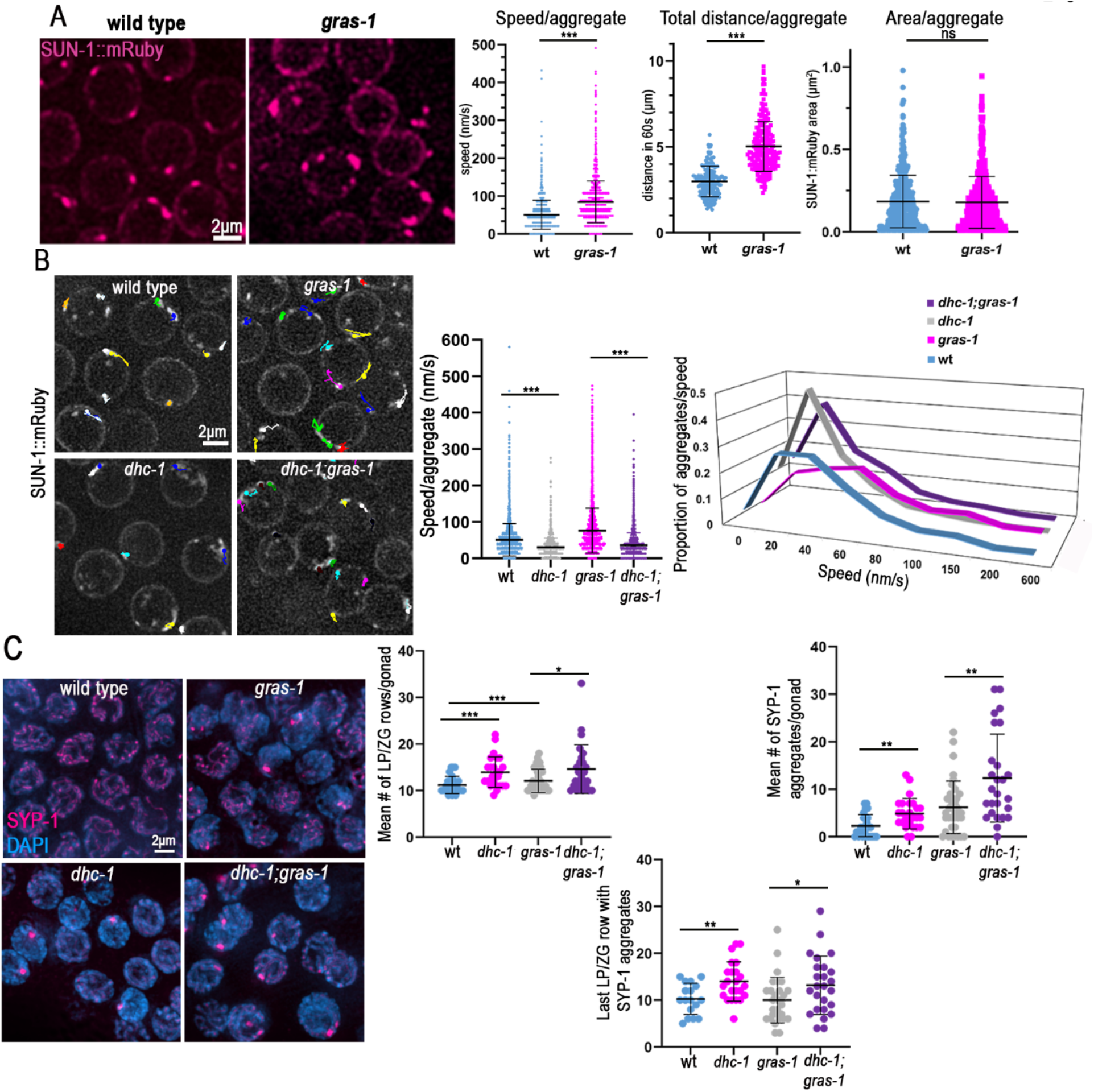
Chromosome movement is limited by GRAS-1 in a dynein-dependent manner during early prophase I. **(A)** Left, snapshots of SUN-1::mRuby live imaging signal (magenta) in wild type and *gras-1* leptotene/zygotene nuclei. Right, dot plots showing the speed (nm/s) of SUN-1::mRuby aggregates, their total distance (μm) traveled in 60s, and their area (μm^2^). ***p<0.001, ns: not significant, by Student’s t-test. n= 173 and 220 aggregates for wild type and *gras-1* for speed and distance and n=529 and 617 for the area measurement, respectively, from two independent biological repeats. **(B)** Left, snapshots from live imaging of SUN-1::mRuby aggregates showing the paths they travelled in 60s in wild type, *gras-1, dhc-1* and *dhc-1;gras-1* leptotene/zygotene nuclei. Right, dot plot displaying the speed (nm/s) of SUN-1::mRuby aggregates and the distribution graph of the aggregates per speed per genotype. ***p<0.0001, Student’s t-test, n= 325, 326, 244, and 377 aggregates, respectively, from between 10 to 13 gonads each, from four independent biological repeats. **(C)** Left, high-resolution images of early pachytene nuclei from wild type, *gras-1, dhc-1*, and *dhc-1;gras-1* co-stained with anti-SYP-1 (magenta) and DAPI (blue). Right top, dot plots displaying the number of rows with nuclei at the leptotene/zygotene (LP/ZG) stage per gonad and the mean number of nuclei with SYP-1 aggregates per gonad (n=26, 25, 36 and 26 gonads, respectively). Right bottom, dot plot showing the last row of nuclei with SYP-1 aggregates per gonad (n=17, 23, 29 and 25, respectively). *p<0.05, **p<0.01, ***p<0.001 by the Mann-Whitney U-test. Bars show the mean and standard deviation in all graphs.

Since the key motor protein involved in promoting early prophase I chromosome movement in *C. elegans* is dynein (Wynne et al. 2012), we examined if the increased SUN-1 speed in *gras-1* mutants was mediated by dynein. Wild type worms depleted of *dhc-1* by RNAi (Fig. S3C; Labrador et al. 2013) exhibited minimal SUN-1 movement with short tracks after 1 minute of imaging and reduced average speed per aggregate (51.37±0.74 nm/s for wild type and 32.66±0.58 for *dhc-1(RNAi)*, p<0.0001, Student’s t-test) (Fig. 3B, video S1). The increased speed of SUN-1 observed in *gras-1;EV* (empty vector) worms was lost in *dhc-1(RNAi);gras-1* worms (76.37±1.03 for *gras-1* and 36.21±0.51 for *dhc-1(RNAi);gras-1*, p<0.0001) (Fig. 3B, video S1). Furthermore, the two types of chromosome movement speeds described for *C. elegans* leptotene/zygotene stage nuclei (processive-chromosome motions with higher speeds in one direction and short-distance movements close to one point; Wynne et al. 2012; Labrador et al. 2013) observed in wild type and exacerbated in *gras-1* were absent upon *dhc-1* depletion with the majority of aggregates displaying a speed around 20-30nm/s (Fig. 3B, rightmost panel). Therefore, the increased speed found in *gras-1* was completely dependent on DHC-1.

Meiotic progression was further impaired in *dhc-1;gras-1* compared to the single *dhc-1* mutant. Compared to wild type, lack of DHC-1 causes a mild extension of the leptotene/zygotene stages (11.19±0.36 and 12.08±0.41 rows of nuclei in wild type and *dhc-1*, respectively, p=0.18, Mann-Whitney U-test) and a significant increase in the presence of SC aggregates in early pachytene (2.27±0.46 and 6.19±0.91 nuclei with SYP-1 aggregates per gonad in wild type and *dhc-1*, respectively, p=0.0044) (Fig. 3C and Sato et al. 2009). *dhc-1;gras-1* double mutant germlines showed a further increase in the number of nuclei with chromatin exhibiting a leptotene/zygotene stage appearance compared to *dhc-1* alone (14.62±1 leptotene/zygotene rows in *dhc-1;gras-1*, p=0.0251) (Fig. 3C), and significantly higher levels of SYP-1 aggregates compared to single mutants (12.35±1.78 nuclei with SYP-1 aggregates in *dhc-1;gras-1* compared to 6.19±0.91 in *dhc-1* and 4.88±0.63 in *gras-1*, p=0.0044 and 0.0005, respectively) (Fig. 3C). Additionally, more of these persistent leptotene/zygotene-like nuclei with SYP-1 aggregates were detected in later stages of prophase I in *dhc-1;gras-1* than in *dhc-1* mutants (13.2±1.22 rows after leptotene/zygotene entry in *dhc-1;gras-1* compared to 10±0.89 in *dhc-1*, p=0.0350) (Fig. 3C, lower panel). Altogether, these data indicate a DHC-1-dependent role for GRAS-1 in limiting chromosome movement/speed in early prophase. However, exacerbated phenotypes, such as the increased number of SYP-1 aggregates, contrast with the epistatic relationship observed for DHC-1 and GRAS-1 in chromosome movement speed, and suggest that GRAS-1 might exert additional functions in regulating meiotic progression.

### GRAS-1 contributes to normal meiotic DSB repair progression

Mutants with altered SC assembly frequently exhibit impaired recombination since the SC is required for normal DSB repair progression and crossover formation (Colaiácovo et al. 2003; Smolikov et al. 2007b, 2009; Bowman et al. 2019). To assess DSB repair in the absence of GRAS-1, we immunostained gonads for RAD-51, a protein involved in strand invasion/exchange steps during homologous recombination (Sung 1994; Alpi et al. 2003; Colaiácovo et al. 2003). *gras-1* mutants exhibited a reduction in the number of RAD-51 foci observed per nucleus from leptotene/zygotene through mid-pachytene stages and a slight increase in late pachytene compared to wild type (p=0.049 for leptotene/zygotene, p<0.0001 for mid-pachytene, and p=0.039 for late pachytene, Mann-Whitney U-test) (Fig. 4A). The increased RAD-51 foci were dependent on the topoisomerase-like SPO-11 protein required for meiotic DSB formation (Fig. S3D). Unrepaired recombination intermediates persisting into late pachytene can result in increased germ cell apoptosis (Gartner et al. 2000). We detected a significant increase in germ cell apoptosis in *gras-1* mutants compared to wild type (3.69±0.21 and 1.97±0.17 mean number of germ cell corpses respectively, p<0.0001, Mann-Whitney U-test) (Fig. 4B). Moreover, the increase in germ cell apoptosis was also meiotic DSB-dependent given that apoptosis levels were no longer elevated in *gras-1* mutants in the absence of SPO-11 (Fig. S3E). Crossover designation levels were not altered as determined by quantification of the number of foci for ZHP-3, the ortholog of budding yeast Zip3 that marks sites designated for crossover formation in late pachytene nuclei (5.99±0.02 and 6.07±0.05 ZHP-3 foci per nucleus in wild type and *gras-1*, respectively, p=0.06, Mann-Whitney U-test). However, a delay in the restriction of ZHP-3 signal from tracks to foci was observed in *gras-1* mutants (Fig. 4C). Analysis of oocytes at diakinesis revealed 6 bivalents in both wild type and *gras-1* mutants with only one oocyte exhibiting a fragile connection between a pair of homologs in *gras-1* (Fig. 4D) (Saito et al. 2009). However, analysis of *gras-1* mutants also lacking the *ced-3* caspase (Yuan et al. 1993), which prevents germ cell apoptosis in late pachytene, revealed an increase in the total number of oocytes with chromosome abnormalities (p=0.0436 compared to wild type, Fisher’s exact test) including univalents, fragile connections, and interbivalent attachments (Fig. 4D). This was accompanied by higher levels of embryonic lethality, larval lethality, and male progeny in *gras-1;ced-3* mutants compared to *ced-3* alone (14.3% vs 5.1% embryonic lethality, p<0.0001; 5% vs 2.5% larval lethality, p<0.0001; and 0.6% vs 0.2% males, p=0.004, Fisher’s exact test) (Fig. 2A). These combined data suggest that GRAS-1 is required for normal meiotic DSB repair progression and maintenance of genomic integrity in the germline.

**Fig. 4.**
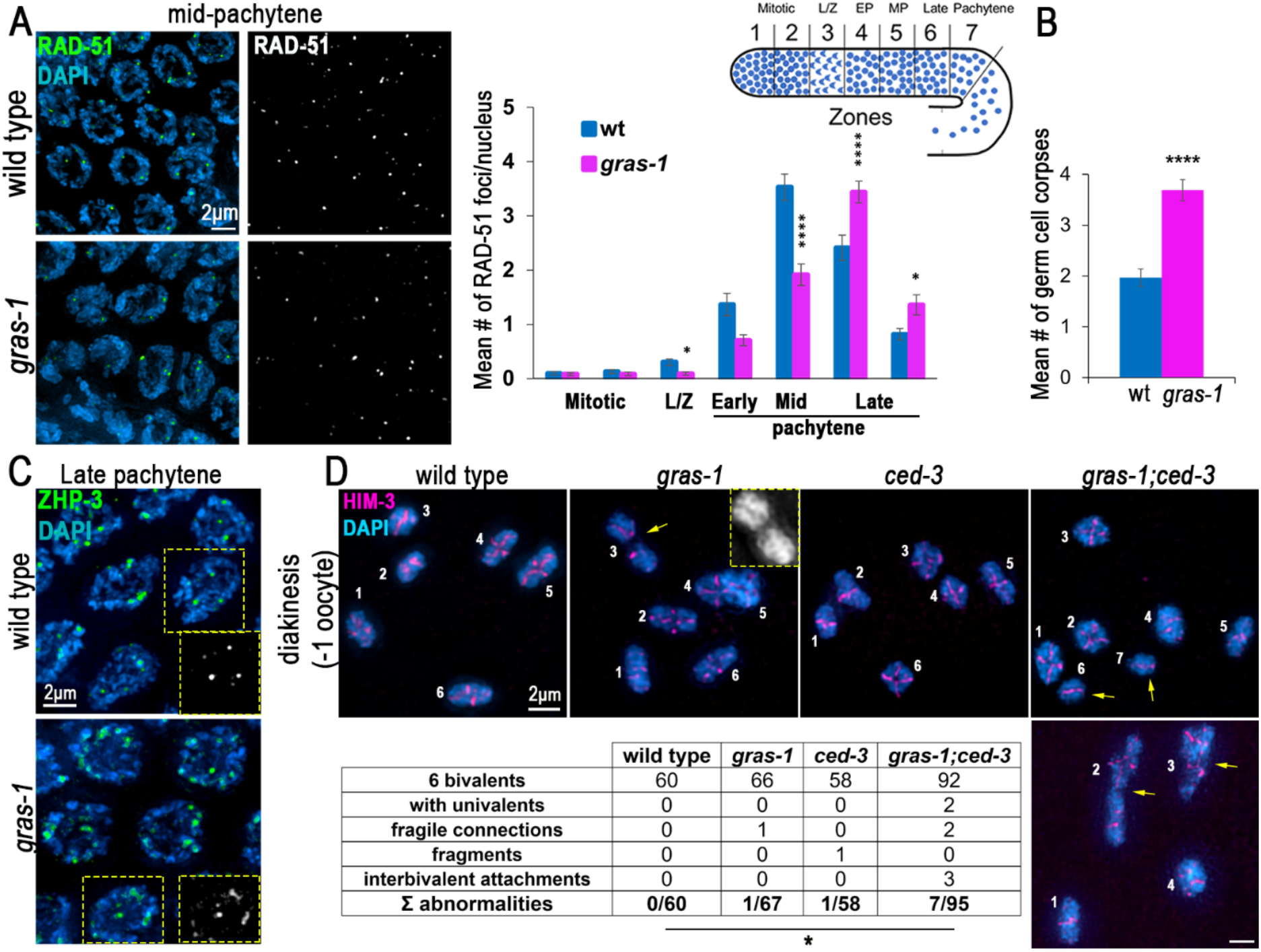
GRAS-1 contributes to normal meiotic DSB repair progression. **(A)** Left, high-resolution images of wild type and *gras-1* mid-pachytene nuclei co-stained with anti-RAD-51 (green) and DAPI (blue). Right, Histogram showing the mean number of RAD-51 foci/nucleus scored along the indicated zones in the germlines of wild type and *gras-1* worms. 3 gonads were scored per genotype and >66 nuclei per zone in two independent biological replicates. Error bars represent the SEM. *p<0.05, ****p<0.0001 by the Mann-Whitney U-test. **(B)** Histogram showing the mean number of germ cell corpses detected in wild type and *gras-1* worms. Error bars represent the SEM. ****p<0.0001, Mann-Whitney U-test, n= 117 and 93 gonads respectively. **(C)** High-resolution images of wild type and *gras-1* late pachytene nuclei stained for ZHP-3 (green) and DAPI (blue). Yellow dotted insets depict ZHP-3 signal in black and white for one of the nuclei in the field showing individual foci in wild type and some ZHP-3 tracks in *gras-1*. n=72 and 71 nuclei each from 15 gonads each. **(D)** Top, High-resolution representative images of diakinesis nuclei stained for HIM-3 (magenta) and with DAPI (blue) from wild type, *gras-1* (yellow arrow indicates a fragile connection), *ced-3*, and *gras-1;ced-3* (yellow arrows indicate univalents in the top and interbivalent attachments in the bottom image). Each individual DAPI-stained body is indicated with a white number. Bottom, table showing the distribution of diakinesis nuclei per genotype that had the normal 6 bivalents or one of the listed abnormalities. *p = 0.0436 between *gras-1;ced-3* and wild type by Fisher’s exact test.

### GRAS-1’s function in limiting chromosome movement in early prophase I is regulated by phosphorylation at a C-terminal S/T cluster

Analysis with different protein phosphorylation prediction programs (Kinase 2.0, NetPhos 3.1 and PHOSIDA) identified S233 and S236 as putative phosphorylation sites within a S/T cluster domain (SSTS) at the C-terminus of GRAS-1 (Fig. 1A). These sites are conserved in human CYTIP (S269 and S270) and GRASP/Tamalin (S293), the former being strongly conserved in other vertebrates (PER viewer) (Pérez-Palma et al. 2020). *In vivo* phosphorylation of GRAS-1 at this S/T cluster was confirmed by mass spectrometry analysis (Fig. 5A and 5B; shown is phosphorylation at S233). Since more than one residue at the SSTS cluster may be phosphorylated, we used CRISPR-Cas9 to edit all four amino acids to either alanine (A) or aspartic acid (D) to generate phosphodead (*gras-1PD*) and phosphomimetic (*gras-1PM*) mutants, respectively (Fig. 5C). Analysis of *gras-1PD* and *gras-1PM* mutants revealed a normal number of eggs laid but increased embryonic and larval lethality compared to wild type (Fig. 5D). *gras-1PD* mutants exhibited levels similar to *gras-1* null (3.1% embryonic lethality in both, p>0.99, and 0.6% larval lethality in *gras-1PD* and 0.97% in *gras-1*, p=0.031, Fisher’s exact test). In contrast, the number of male progeny in phosphodead or phosphomimetic mutants was indistinguishable from wild type (Fig. 5D, right panel). To assess whether GRAS-1 phosphorylation is required for its role in limiting chromosome movement in early prophase I, we analyzed the speed of SUN-1::mRuby aggregates in *gras-1PD* and *gras-1PM* mutants (Fig. 5E, videos S2). The inactivation of the phosphorylation domain in *gras-1PD* produced a higher average speed per aggregate compared to wild type (65.18±1.12 nm/s and 51.37±0.74 nm/s, respectively, 325 and 245 aggregates, p <0.0001, Student’s t-test), but not as elevated as in the *gras-1* null mutant (73.17±1.09 nm/s, 270 aggregates, p<0.0001). In contrast, mimicking a phosphorylated SSTS domain resulted in chromosome movement speeds similar to those observed in wild type (53.05±0.65 nm/s in *gras-1PM*, 343 aggregates, p=0.0872). Depletion of dynein by RNAi in the phosphodead and phosphomimetic mutants resulted in SUN-1 aggregates indicating non-processive chromosome motions with slower speeds (Fig. 5E, lower right panel). These results suggest that GRAS-1 function is regulated by phosphorylation at these conserved C-terminal residues.

**Fig. 5.**
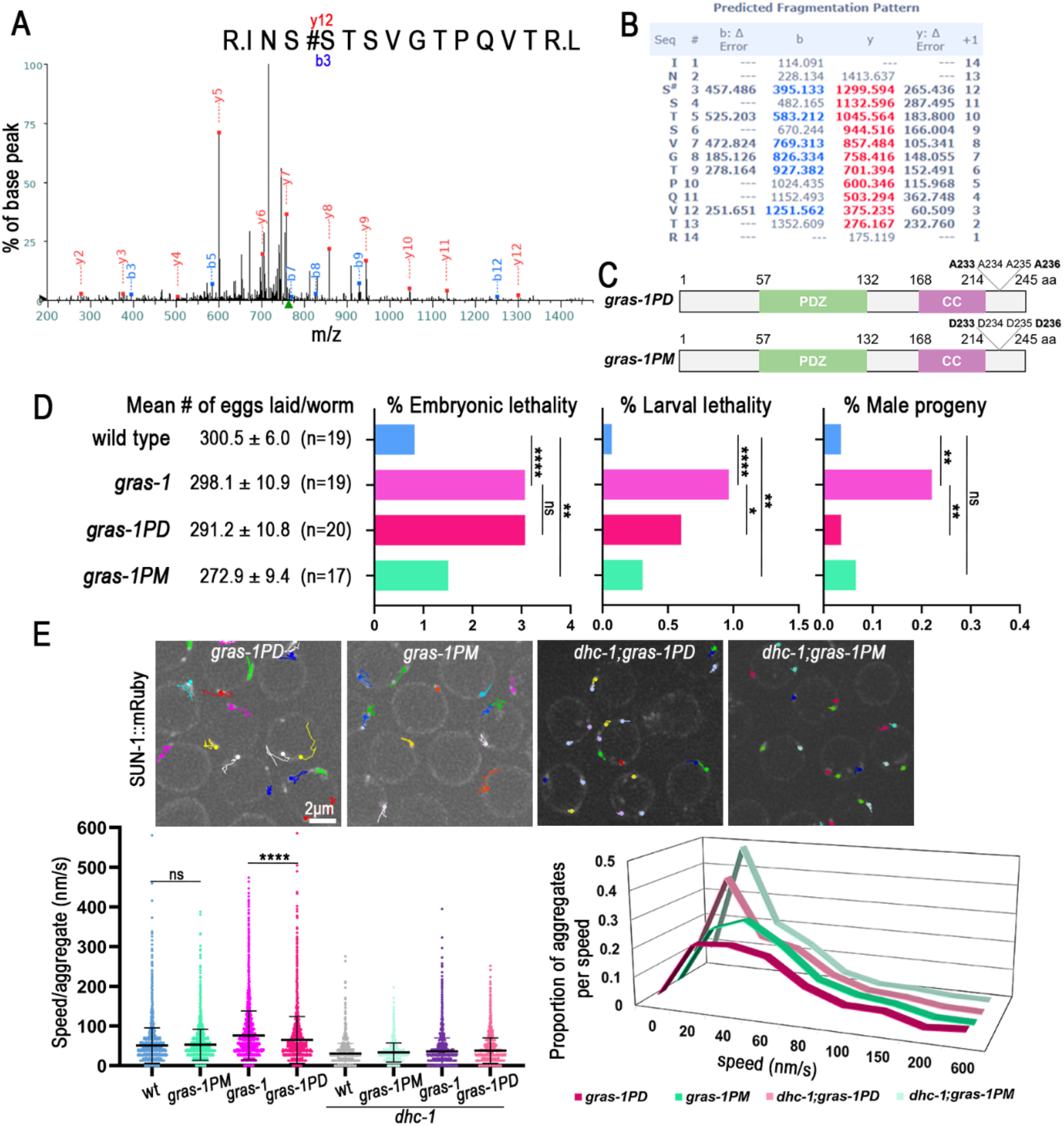
GRAS-1 function in limiting early prophase I chromosome movement is regulated by phosphorylation. **(A)** Mass spectrometry fragmentation spectrum for GRAS-1::GFP peptide INSSTSVGTPQVTRL in the range 200-1400 m/z. The annotated spectrum shows fragment ion species matched between theoretical and measured values. “b-ions” are generated through fragmentation of the N-terminal peptide bond, and “y-ions” through the C-terminal. **(B)** Predicted MS fragmentation pattern and deviations (∆ Error). Analysis of b and y ions is consistent with the phosphorylation of the second serine (S#) in this peptide, corresponding to S233 of the GRAS-1 protein. **(C)** Schematic representation of the proteins encoded by the CRISPR-Cas9 engineered *gras-1PD* (phosphodead) and *gras-1PM* (phosphomimetic) mutants. A: alanine, D: aspartic acid. **(D)** Shown are the mean number of eggs laid (brood size) ± SEM, percentage of embryonic lethality, larval lethality, and males for the indicated genotypes. *p < 0.05, **p < 0.01, ****p < 0.0001, ns: not significant, by Fisher’s exact test. n= number of worms for which entire progeny were analyzed. **(E)** Top, snapshot of live imaging of SUN-1::mRuby aggregates and their travelled paths in 60s in *gras-1PD, gras-1PM, dhc-1;gras-1PD* and *dhc-1;gras-1PM* leptotene/zygotene nuclei. Bottom, dot plot displaying the speed (nm/s) of SUN-1::mRuby aggregates and the distribution graph of the aggregates per speed for the indicated genotypes. Worms were grown in bacteria containing the empty-vector or *dhc-1(RNAi)* construct. ***p<0.0001, ns: not significant by Student’s t-test, n= 325, 343, 326, 245, 244, 195, 377 and 247 aggregates per genotype as shown in figure, from 9 to 13 gonads and at least two independent biological repeats. Error bars represent the mean ± SD.

### GRAS-1 shares partial functional conservation with human CYTIP

The fact that GRAS-1 protein structure, phosphorylation, and reproductive tissue expression seem to be conserved in mammals (Fig. 1A, S1A) suggests that similar functions could be performed by either CYTIP and GRASP/Tamalin in mammals. To test this possibility, we first examined *Tamalin, Cytip* DKO mouse mutants (Fig. S4A). The mouse mutants had fertility rates, testis weight, and seminiferous tubule morphology equivalent to littermate controls (Fig. S4B). Analysis of meiotic progression in chromatin spreads from both male and female *Tamalin, Cytip* DKOs immunostained with γH2AX to assess DSB formation and SYCP3 to examine chromosome synapsis did not reveal any defects compared to controls in oocytes (Fig. 6A) and spermatocytes (Fig. S4C). Analysis of DSB repair progression by immunostaining meiotic prophase I cells for RPA revealed normal levels relative to controls in oocytes (Fig. 6B), and the ATR DNA damage response kinase and RPA in spermatocytes (Fig. S4D, E). The formation of the central element of the SC also showed normal progression in oocytes (Fig. S5A). Finally, the number of crossover recombination events determined by assessing MLH1 foci in spermatocytes and CDK2 in oocytes from *Tamalin, Cytip* DKO mice was the same as in wild type (Fig. S5B, 1.09 ± 0.02, n=409, and 1.08 ± 0.02, n=416, MLH1 foci on chromosomes in the control and DKO, respectively, p=0.7641 Mann-Whitney test; Fig. S5C, 3.14 ± 0.03, n=427, and 3.18 ± 0.02, n=451, CDK2 foci in the control and DKO respectively, p=0.2332). These results are similar to the crossover analysis in the worm *gras-1* mutant in which levels of ZHP-3 foci were indistinguishable from wild type.

**Fig. 6.**
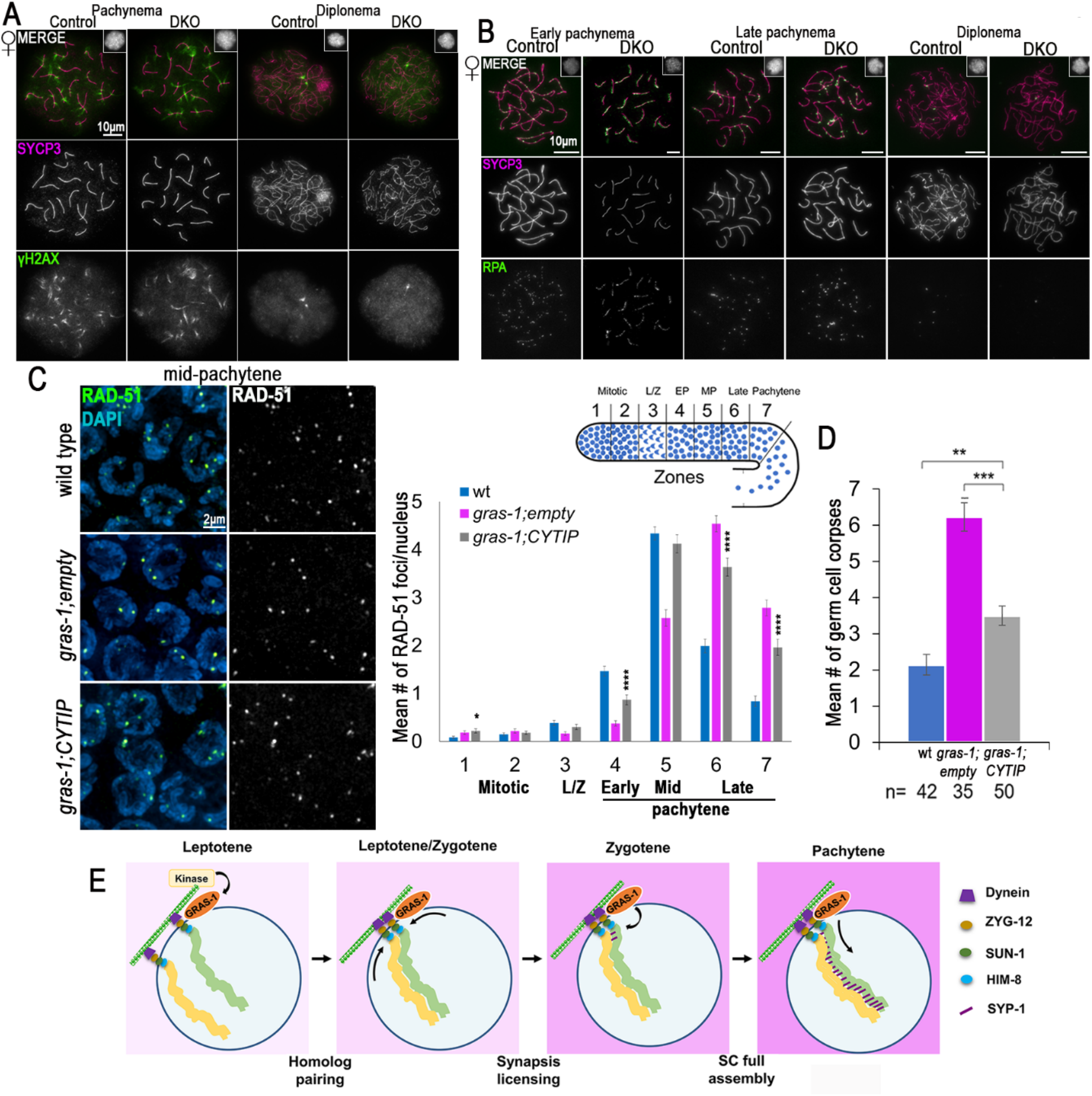
GRAS-1 shares partial functional conservation with human CYTIP. **(A)** Chromatin spreads from mid meiotic prophase (pachynema) and late meiotic prophase (diplonema) control and *Tamalin-Cytip* DKO *Mus musculus* oocytes co-immunostained with anti-SYCP3 (magenta) and anti–γ-H2AX (green). Insets show normal chromatin morphology (DAPI). n=50 cells per mouse and 3 mice per genotype. **(B)** Chromatin spreads from early and late pachynema and diplonema, control and *Tamalin-Cytip* DKO *Mus musculus* oocytes co-immunostained with antibodies against SYCP3 (magenta) and RPA (green). Insets show normal chromatin morphology (DAPI). **(C)** Left, high-resolution images of mid-pachytene nuclei from wild type, *gras-1;empty* and *gras-1;CYTIP* germlines stained with anti-RAD-51 (green) and DAPI (blue). Right, Histogram showing the mean number of RAD-51 foci/nucleus scored along the germlines for the indicated genotypes. 5-6 gonads were scored per genotype in two independent biological replicates. Error bars represent the SEM. *p<0.05, ****p<0.0001 by the Mann-Whitney U-test. **(D)** Histogram showing the mean number of germ cell corpses of wild type (*gras-1;empty* and *gras-1;CYTIP* worms. Error bars represent the SEM. **p<0.01, ***p<0.001, Mann-Whitney U-test, n= 42, 35 and 50 gonads, respectively. **(E)** A model for the role of GRAS-1 during *C. elegans* meiosis. We propose that GRAS-1 bridges the cytoskeleton, the LINC complexes, and chromosomes to limit chromosome movement in a phosphorylation-dependent manner and license synapsis during early prophase I. A single pair of homologous chromosomes (yellow and green) is shown for simplicity within a nucleus delimited by the nuclear envelope (dark blue line) and attached to the LINC complex and a single microtubule (green/white checkered bar).

Even though mutant *Tamalin, Cytip* DKO mice did not present obvious fertility or meiotic defects, GRAS-1 function could still be conserved to a lesser extent or diverged in mice but not in other vertebrates. To assess functional conservation with the human orthologs, we complemented *gras-1* null worms with the human cDNA of CYTIP, the ortholog with the highest sequence similarity. Using the SKI LODGE system (Silva-García et al. 2019) we introduced a cassette into chromosome III with expression of the human coding sequence driven by the *pie-1* germline-specific promoter (Fig. S5D). We examined DSB repair progression by RAD-51 immunostaining of germlines from wild type, *gras-1* null carrying an empty vector cassette inserted in chromosome III, and *gras-1* expressing human CYTIP (Fig. 6C). *gras-1;CYTIP* exhibited a partial rescue relative to *gras-1* with RAD-51 levels increasing in early pachytene, albeit not reaching the same levels as in wild type until mid-pachytene (1.47±0.10 foci/nucleus in wild type and 0.87±0.10 in *gras-1;CYTIP*, p<0.0001, Mann-Whitney U-test), and a partial reduction in the levels observed in late pachytene (1.99±0.14 and 0.84±0.11 foci in late pachytene in zones 6 and 7, respectively, in wild type; 3.64±0.19 and 1.96±0.17 in *gras-1;CYTIP*, p<0.0001). Therefore, human CYTIP expression in *gras-1* null mutants resulted in an intermediate phenotype between *gras-1* null and wild type. Similarly, we observed reduced levels of germ cell corpses in *gras-1* worms complemented with CYTIP compared to *gras-1* null (6.23±0.40 in *gras-1;empty* and 3.5±0.27 in *gras-1;CYTIP*, p<0.0001, Fisher’s exact test), but not a complete reversion to wild type levels (2.14±0.28, p=0.001) (Fig. 6D). A partial rescue of the SC assembly defects was also observed with lower levels of SC aggregates in early pachytene (Fig. S5E). These results suggest that GRAS-1 protein function could play similar roles in vertebrates, but the divergence of the proteins and the duplication might affect the processes involved.

## DISCUSSION

Early prophase I events are key determinants of correct chromosome segregation at meiosis I and therefore need to be tightly coordinated. In the present study, we uncover a layer of regulation for these events mediated by the conserved GRAS-1 protein. We propose that GRAS-1 connects the stabilization of homologous chromosome pairing with the licensing of SC formation between homologs by limiting chromosome movement during early prophase I (Fig. 6E). We suggest that GRAS-1 acts as a scaffold connecting proteins implicated in both processes, thereby establishing a physical link between the nuclear envelope and cytoplasmic structures.

GRAS-1 expression increases in germline nuclei during the transition from mitosis into meiotic prophase I (Fig. 1C, S1B) and GRAS-1 localizes near the germ cell NE (Fig. 1E, S1D). GRAS-1 localization seems to be more dispersed compared to the membrane actin fibers and cell membrane components such as SYX-4 (Sato et al. 2008) (Fig. S1D). GRAS-1 contacts the NE cytologically (Fig. 1D, E) and mass spectrometry results from GRAS-1::GFP pull-downs show that GRAS-1 may interact directly or indirectly with several NE proteins and numerous tubulin, actin, spindle, and chromosome segregation proteins (Fig. 1F, G). Among these are separase, a caspase-related protease that regulates sister chromatid separation (Alexandru et al. 2001; Hornig and Uhlmann 2004), the protein phosphatase PP1 orthologs GSP-1/2 with various roles including regulation of sister chromatid cohesion upon entrance into meiosis (Ceulemans and Bollen 2004; Tzur et al. 2012), and PLK-1, whose meiotic role in phosphorylating key chromosome movement regulators and SC components could be an important effector for GRAS-1 function during early prophase I (Labella et al. 2011; Woglar et al. 2013; Nadarajan et al. 2017). Another putative interaction partner of GRAS-1 is LMN-1, which also plays an important role in chromosome movement and supports a connection for GRAS-1 with structural components of the NE (Phillips et al. 2005; Link et al. 2018). The motor protein kinesin KLP-17 provides an additional target by which GRAS-1 chromosome movement functions could be connected. Kinesins produce opposite movements of cargo proteins to dyneins and the *C. elegans*-specific KLP-17 protein is expressed in the germline, has microtubule binding activity, and has been proposed to have chromosome movement and segregation activity (Siddiqui 2002; Robin et al. 2005; Heppert et al. 2018). The KASH domain protein KDP-1 is implicated in cell cycle progression and its localization depends on SUN-1 in the germline (McGee et al. 2009). The LINC complex protein KDP-1 interacts with SUN-1 or UNC-84 at the NE and could interact with other SUN-1 partners, although its role during meiosis requires further investigation. Finally, nucleoporins and importins identified in the pull-downs (NPP-3, NPP-13, IMA-2, and IMA-3) have been implicated in regulating chromosome attachment to the NE, chromosome movement, meiotic recombination, chromosome segregation, and the timely incorporation of SC proteins (Bowman et al. 2019; Palacios et al. 2021). However, our analysis of *gras-1* in combination with *ima-1* and *ima-2* mutants does not support them acting in the same pathway.

Our data indicate that GRAS-1 acts to limit chromosome movement (Fig. 3). GRAS-1 could impose resistance to the free movement of chromosomes from outside of the nucleus when they find a homologous partner, thereby stabilizing that connection (Fig. 6E). In *C. elegans*, similarly to mice and *S. pombe*, the cytoskeletal forces driving the movement of chromosomes from outside the nucleus are controlled by microtubules and the motor protein dynein connecting to the chromosome-LINC complex (Sato et al. 2009; Wynne et al. 2012; Zetka et al. 2020). GRAS-1 may function in the same pathway and limit the action of dynein and microtubules since dynein depletion in the absence of GRAS-1 results in chromosome movements similar to those in the dynein mutant alone (Fig. 3B). Although mutations in co-chaperone FKB-6 increase meiotic chromosome movement (Alleva et al. 2017), GRAS-1 must act through a different pathway since *fkb-6* mutants showed decreased resting time between chromosome movements, whereas aggregates in *gras-1* had increased general speeds (Fig. 3A). Further, the *fkb-6* mutant in combination with either *dhc-1* depletion or a *zyg-12* mutant did not exhibit exacerbated defects in SC formation or chromosome pairing, in contrast with *dhc-1(RNAi);gras-1* mutants where these defects are accentuated (Fig. 3C). Moreover, FKB-6 was not identified in GRAS-1::GFP pull-downs, and its localization was more dispersed throughout the cytoplasm in contrast to the membrane localization for GRAS-1. Further, FKB-6 expression was not meiosis-specific, which is connected with a role for FKB-6 in regulating microtubule formation and mitotic segregation in the *C. elegans* germline (Alleva et al. 2017). Additionally, cytoplasmatic protein vinculin/DEB-1 has also been proposed to limit the movement of LINC complexes and produce abnormal synapsis (Rohožková et al. 2019). We believe GRAS-1 functions in a different way than vinculin/DEB-1 because of the differences in localization and phenotypes: *deb-1* mutants had a high number of univalents at diakinesis, defects in the loading of proteins along meiotic chromosome axes (which could be the cause of the severe synapsis defects observed), and their pairing defects are opposite to that in *gras-1* since they initially have the same level of pairing as wild type worms, but then homologs do not achieve complete pairing in most pachytene nuclei.

The excess chromosome movement found in the absence of GRAS-1 could be the reason for the extension in leptotene/zygotene stages, the pairing delays, and the altered DSB repair progression observed in the germline (Fig. 2B, C and Fig. 4). However, GRAS-1 could be involved in transmitting additional signals once homologs find a partner, since in *dhc-1;gras-1* double mutants there were more instances of SC aggregates and those appear in late pachytene (Fig. 3C and Fig. 6E). One possibility is that GRAS-1 helps license the initial assembly of the SC from the PC ends of paired chromosomes, so that in the absence of GRAS-1 homologs do not stay together long enough and the imported SC proteins self-aggregate. However, if that were the case we would expect the SC defects to affect most nuclei, as observed for the defects in chromosome movement. Alternatively, GRAS-1 may regulate the loading of SC proteins via a yet unknown mechanism, given the incomplete polymerization of SYP-1 observed in *gras-1* mutants at mid-pachytene stage (Fig. 2D, E). This is further supported by the presence of SYP-3 in GRAS-1::GFP pull-downs assessed on westerns and interactions with SYP-1/2/3 in yeast two-hybrid assays (Fig. S3A, B). If GRAS-1 regulates SC assembly and/or loading, it does so in a manner that is distinct from the combined role of Akirin with importins (Bowman et al. 2019) since we did not find evidence of cohesin or axial element defects in *gras-1* mutants alone, or in combination with *ima-1* and *ima-2* (Fig. 1E, S2C).

GRAS-1 is conserved in animals, and the gene underwent a duplication event in chordates resulting in CYTIP and GRASP/Tamalin (Fig. 1A, B). All three proteins carry PDZ and coiled-coil domains, usually involved in protein-protein interactions. In addition, they carry a disorganized C-terminal region (longer in the mammalian orthologs) that could be involved in regulating their function since a phosphorylated serine in the S/T cluster is conserved in both human CYTIP and GRASP. In addition, GRAS-1 protein interactions might also be conserved since mammalian CYTIP and GRASP have been found to interact with Cytohesin-1 (Heufler et al. 2008; Kitano et al. 2002; Teuliere et al. 2014) and the worm ortholog, GRP-1, was a top and specific hit in all GRAS-1::GFP pull-down experiments. Similar to worm *gras-1* mutants, *Tamalin, Cytip* DKO mice did not exhibit severe fertility defects or crossover recombination problems (Fig. 6). However, DKO mice also did not show defects in meiotic progression, chromosome synapsis, and DSB repair progression (Fig. 6, S4, S5). Meiotic progression defects may be more easily detected in *gras-1* mutant worms because of the spatiotemporal organization of meiosis within intact worm gonads that facilitates the observation and quantification of subtle defects compared to individual cell spreading techniques in mouse samples. Gene duplication divergence might also explain these differences because CYTIP and GRASP are sometimes expressed in different tissues and often have distinct roles (Uhlén et al. 2015; Yanpallewar et al. 2012; Coppola et al. 2006). However, protein structure and functions could still be conserved throughout evolution to partially complement worm GRAS-1 function with the closest human ortholog, CYTIP (Fig. 6 and S5D, E). Moreover, there could be differences between the mouse and human proteins, or subtle phenotypes or timing issues in the double KO that we could not detect.

In conclusion, we propose a model for the conserved GRAS-1 protein during meiosis in which its localization and protein interactions limit the movement of chromosomes in early prophase I. GRAS-1 may function to stabilize connections between homologs by serving as a protein scaffold connecting the NE environment with cytoskeletal forces to license SC assembly (Fig. 6E).

## MATERIALS AND METHODS

### Worm strains and growth conditions

N2 Bristol worms were used at the wild-type background. Lines were cultured under standard conditions as in (Brenner 1974). Some mutant lines were obtained from the Caenorhabditis Genetics Center (CGC) and from the National BioResource Project for the nematode *C. elegans* (NBRP, Japan). *gras-1* mutant lines were generated using the CRISPR-Cas9 system (Kim and Colaiácovo 2016; Tzur et al. 2013). A deletion from +295 to 189 post termination codon nucleotides was initially generated (*rj15*). Full deletion lines from −28 to 52 post termination codon nucleotides (*rj27*) and start codon to 27 nucleotides after the stop codon (*rj28*), not affecting the promoter of operon CEOP1424, were generated using sgRNA GTTTATCTCTGAACACTCAT and the PAM sequence was mutated from GGG to AGA. The *gras-1(rj28)* allele was used for these studies since the deletion in *rj27* partly extends into the promoter.

A *gras-1::gfp* line was generated using sgRNA TACTAGAGACGCGTGACTTG, a linker (ggcggcagcggc) and GFP sequence (pPD95.67) before the stop codon. The guideRNA sequence was mutated to avoid re-cutting. Phosphodead (*gras-1PD*) and phosphomimetic (*gras-1PM*) mutants were produced using the sgRNAs CACGCTTTACGAACTTGAT and TTTACTAGAGACGCGTGACT, respectively, and by changing the PAMs or sgRNA region to synonymous codons to avoid re-cutting. Changes in codons 233 to 236 were made so that SSTS amino acids were mutated into AAAA or DDDD, respectively. All three CRISPR-Cas9-engineered lines were produced by SunyBiotech (Fu Jian, China).

Complementation lines expressing HsCYTIP (Dharmacon, MHS6278-202807568) and HsGRASP (Dharmacon, MHS6278-202759705) cDNAs were generated using the SKI-LODGE system (Silva-García et al. 2019). HsCYTIP was inserted into *wbmls60[pie-1p::3xFLAG::dpy-10 crRNA::unc-54 3’UTR, III]* using *dpy-10* crRNA as a target and a PCR template with homology arms including a GFP artificial intron (pPD95.67, gtaagtttaaacatatatatactaactaaccctgattatttaaattttcag) before the CYTIP start codon. CRISPR-generated mutations were Sanger-sequenced (Macrogen). CGC and NBRP mutants were outcrossed at least 6 times with N2. CRISPR-Cas9 generated lines were outcrossed at least 4 times with N2. A full list of the strains used in this study can be found in Supplemental Table 1.

### Yeast two-hybrid analysis

GRAS-1 was found in a yeast two-hybrid screen designed to identify proteins interacting with the SC central region protein SYP-3 (Smolikov et al. 2009). This was confirmed by using GRAS-1 full length, ∆Nt^69-245^, and ∆Ct^1-163^ cloned into Gateway destination vector pVV213 (activation domain, AD). pVV212 (Gal4 DNA binding domain, DB) was used to clone SYP-1/2/3/4 full length, N-terminal, and C-terminal constructs described in (Smolikov et al. 2009; Schild-Prüfert et al. 2011). Strains were mated on YPD and selected on SC Leu- Trp- plates as described in (Walhout and Vidal 2001; Saito et al. 2012).

### Immunoprecipitation and MS analysis

24h post-L4 worms expressing GRAS-1::GFP were collected, frozen, and homogenized, and an anti-GFP antibody used for immunoprecipitation as in (Nadarajan et al., 2016 and Gao et al., 2015), in four independent experiments. To identify the interacting proteins in GRAS-1::GFP pull-downs and examine the phosphorylation status of GRAS-1, a proteoExtract protein precipitation kit (Calbiochem, #539180) was used followed by mass spectrometry analysis (Taplin Biological Mass Spectrometry Facility, HMS, MA). Protein interactors were curated using 4 independent controls and the 4 independent GRAS-1::GFP experiments using the Normalized Spectral Abundance Factor method (Zybailov et al. 2006), normalizing by protein weight, the total number of peptides per experiment, substituting the hits not found in a particular experiment by the 100 times lowest percentile, and bait correction factor. Fold-change relative to the controls was calculated using the average of the 4 experiments normalized to the bait peptides. T-student test was used to determine which proteins with greater than 1.5-fold-change were statistically significant and corrected by the number of hypothesis. A volcano plot was generated using the log2 of the fold-change and −log10 of the p-value.

### C. elegans immunofluorescence and imaging methods

*C. elegans* gonads from 24 hour post-L4 hermaphrodites were dissected and whole-mounted on slides as in (Colaiácovo et al. 2003) using 1% paraformaldehyde fixation, or 4% for the α-RAD-51 time course analysis. A list of primary antibodies used in this study along with their corresponding dilutions can be found in Supplemental Table 2. Secondary antibodies were purchased from Jackson ImmunoResearch Laboratories (West Grove, PA) as AffiniPure IgG (H+L) with minimum crossreactivity: α-rabbit Cy3, α-goat Cy3 (1:200); α-chicken Alexa 488, α-rabbit Alexa 488, α-guinea pig Alexa 488 (1:500); and α-rabbit Cy5, α-goat Cy5, α-guinea pig Cy5, α-chicken Alexa 647 (1:100).

High-resolution imaging was performed with a IX-70 microscope (Olympus, MA) at 0.2μm Z-intervals usually dividing the gonad in 7 equally sized zones from the distal tip with a cooled CCD camera (CH350; Roper Scientific, AZ) driven by the DeltaVision Imaging System (Applied Precision, GE Healthcare). Fixed samples were imaged using a 100x objective (N.A. 1.4), 10X eyepieces, and an auxiliary magnification lens of 1.5X for imaging diakinesis oocytes. Images were deconvolved using a conservative ratio and 15x cycles with SoftWorx 3.3.6 software from Applied Precision, and processed with Fiji ImageJ (Schindelin et al. 2012).

Super-resolution imaging of 24 hour post-L4 *gras-1::gfp;sun-1::mRuby* worms was performed with an OMX 3D-Structured Illumination microscope with focus drift collection after point-spread function assessment (Nikon Imaging Center, Harvard Medical School).

### Pairing measurements

Quantitative time course analysis of homologous chromosome pairing was assessed by immunostaining 24 hour post-L4 dissected gonads with α-HIM-8. Gonads were divided into seven 512×512 pixel zones, with Zone 1 starting approximately three nuclear diameters away from the distal gonad tip as in (MacQueen 2001). HIM-8 foci were considered paired if ≤0.75μm apart. Two independent biological replicates and a total of 6 gonads were scored for each genotype. The average number of nuclei scored per zone in wild type and *gras-1* was respectively: zone 1 (n=160, 134), zone 2 (n=142, 133), zone 3 (n=144, 150), zone 4 (n=157, 167), zone 5 (n=165, 178), zone 6 (n=136, 165), and zone 7 (n=131, 145).

### RAD-51 and ZHP-3 time course analyses

Whole-mounted gonads from 24 hour post-L4 hermaphrodites immunostained either for RAD-51 or ZHP-3 were divided into seven equal-size zones with two independent biological replicates per comparison. Fiji plugin Cell Counter (https://imagej.nih.gov/ij/plugins/cell-counter.html) was used to track in 3D the number of foci for each nucleus in a zone. The average number of nuclei scored per zone for the RAD-51 analysis was: zone 1 (n=97.8), zone 2 (n=114), zone 3 (n=115), zone 4 (115.6), zone 5 (n=113.4), zone 6 (n=108), and zone 7 (n=101.6). The number of nuclei scored in zones where ZHP-3 signal was restricted to individual foci was 72 (wild type) and 71 (*gras-1*).

### Plate phenotyping

Between 10 to 15 L4-stage hermaphrodites for each genotype were placed on individual NGM plates freshly seeded with *E. coli* OP50 to score the total numbers of eggs laid (brood size), embryonic lethality (number of unhatched eggs/total number of eggs laid), larval lethality (number of dead larvae/total number of hatched eggs) and male frequency (number of males /total number of adult worms). Individual P0 worms were moved every 24 hours to new plates for four consecutive days to score entire brood sizes.

### RNAi by feeding

Feeding RNA interference experiments were performed as in (Govindan et al. 2006) using HT115 bacteria expressing pL4440 empty vector as a control and bacteria expressing dsRNA for the gene of interest from the Ahringer RNAi library (*ima-2* F26B1.3, *dhc-1* T21E12.4) (Source Bioscience). Between three to five L4-stage worms were placed per plate (in a minimum of 2 plates per genotype per replicate) and grown at room temperature. F1 L4 animals were transferred to newly seeded RNAi plates and 24h post-L4 worms were analyzed. Alternatively, P0 L1-stage animals were placed in RNAi plates at 25ºC and analyzed 24h post-L4 stage when performing *dhc-1* depletion experiments.

### Germ cell apoptosis experiments

The number of germ cell corpses per gonad arm was scored in 20h post-L4 stage worms as in (Kelly et al. 2000). A minimum of 30 gonads were scored for each genotype using a Leica DM5000B fluorescence microscope.

### Bioinformatics and databases

The evolutionary tree of GRAS-1 family protein members was obtained from TreeFam (2019 TF316315, Ruan et al., 2007, http://www.treefam.org/). Degree of conservation between CeGRAS-1 and HsCYTIP or HsGRASP was calculated using NCBI-blast (https://blast.ncbi.nlm.nih.gov/Blast.cgi) and NCBI-cobalt (http://www.ncbi.nlm.nih.gov/tools/cobalt/). PDZ domain prediction was performed using the ExPaSy Prosite tool (Sigrist et al., 2012, http://prosite.expasy.org/). ExPaSy-Marcoil tool was used to predict coiled-coil domains (Delorenzi and Speed, 2002, http://bcf.isb-sib.ch/webmarcoil/webmarcoilC1.html). *C. elegans* gene expression was assessed using NEXTDB (nematode.lab.nig.ac.jp/) and (Ortiz et al. 2014; Reinke 2004; Tzur et al. 2018).

### Chromosome movement assessment by live imaging

Wild-type and *gras-1* hermaphrodites carrying the *oxls279[Ppie-1::GFP::H2B, unc +](II); ieSi21 [sun-1p::sun-1::mRuby::sun-1 3'UTR + Cbr-unc-119(+)] IV* constructs were grown at 20ºC or 25ºC and selected at the L4 stage. 14-16h post-L4 live worms were mounted on 2% agarose pads with M9 containing 0.01% levamisole. Hyperstack images (x, y, z, t) at 594nm for SUN-1::mRuby fluorescence, were taken using the 60x or 100X objective at 0.2μm intervals. Nuclei with chromatin in the crescent-shaped configuration characteristic of the leptotene/zygotene stage were imaged every 5 seconds for a minute and SUN-1 aggregate trajectory was followed. An additional stack (x, y, z) capturing GFP::H2B signal at 523nm was collected as a reference for the chromatin shape. Images were registered using the Fiji (NIH) plugin Manual Registration. 2D speed analysis of SUN-1::mRuby aggregates was performed using the Fiji Manual Tracking plugin as in (Alleva et al. 2017; Link et al. 2018).

### Ethics statement

All mice used were bred at the Johns Hopkins University Bloomberg School of Public Health (JHSPH, Baltimore, MD) in accordance with criteria established by the National Institute of Health and the U.S. Department of Agriculture. The Johns Hopkins University Institutional Animal Care and Use Committee (IACUC) approved the protocols for the mice’s care and use.

### Mice mutant lines

*Tamalin* KO mice were kindly provided by Dr. Lino Tessarollo (Yanpallewar et al. 2012). We obtained mice harboring the *Cytip* tm1a “knockout first” allele from the Mutant Mouse Resource and Research Centers (MMRRC) at University of California-Davis. The *Cytip* tm1a allele has loxP sites flanking exon 4 and 5. Mice heterozygous for *Cytip* tm1a were bred with mice harboring the *Spo11-Cre* transgene (C57BL/6-Tg Spo11-cre)Rsw/PecoJ), which express Cre recombinase in spermatocytes shortly after meiotic entry (Lyndaker et al. 2013). The resulting progeny from this cross harbored the *Cytip* tm1b KO allele. We subsequently bred mice harboring the Tamalin KO and Cytip KO alleles to create the *Tamalin*, *Cytip* DKO mice for analysis. We also bred mice heterozygous for *Cytip* tm1a allele with mice harboring FLP recombinase transgene (FLP tg/0) to produce progeny with the *Cytip* tm1c “conditional knockout” (cKO) allele. These mice were used to create the *Cytip* cKO mice that were homozygous for the *Cytip* tm1c allele and hemizygous for the *Spo11-Cre* transgene.

### Mouse genotyping

Mouse genotypes were obtained by polymerase chain reaction (PCR). Mice toe tips were digested in 50 mM NaOH at 95°C for 15 mins and 1M Tris-HCl pH 7.5 was added to the digestion. The digested toe tips were used as the DNA template in the PCR.Primers used in the PCRs are listed in Supplemental Table 3. PCR conditions: 90°C for 2 min, 30 cycles of 90°C for 20 s, 58°C for 30 s, 72°C for 1 min. PCR products were analyzed using 2% agarose gels.

### Histological analysis and tubule squash preparations

Testes were fixed in Bouins fixative, embedded in paraffin, and serial sections of 5-μm thickness were placed onto slides and stained with hematoxylin and eosin (H&E). Mouse tubule squashes were prepared as described in (Wellard et al. 2018).

### Mouse chromatin spread preparations and imaging

Spermatocyte and oocyte chromatin spreads were prepared as previously described (Jordan et al., 2012; Wellard *et al.* 2022; Hwang *et al.* 2018). Primary antibodies and dilution used for immunolabeling are presented in Supplemental Table 4. Secondary antibodies against human, rabbit, rat, mouse, and guinea pig IgG and conjugated to Alexa 350, 488, 568, or 633 (Life Technologies) were used at a 1:500 dilution.

Images from chromatin spread preparations were captured using a Zeiss CellObserver Z1 microscope linked to an ORCA-Flash 4.0 CMOS camera (Hamamatsu). Testis sections stained with H&E staining were captured using a Zeiss AxioImager A2 microscope linked to an AxioCam ERc5s camera, or Keyence BZ-X800 fluorescence microscope. Images were analyzed and processed using ZEN 2012 blue edition imaging software (Zeiss) or with BZ-X800 Viewer and Analyzer software (Keyence).

### Statistical methods

The average of the data was used as a typical representation throughout the manuscript, accompanied by the standard error as a measure of data deviation. Statistical tests were performed in GraphPad Prism 8. Variables with continuous data, such as speed, distance, and area, were compared using unpaired 2-tailed t-tests. The Fisher exact test was used to assess the statistical significance for the distribution of data in the samples. All other comparisons were tested using the two-sided non-parametric Mann Whitney U-test. Graphs for comparisons were generated in Microsoft Excel or GraphPad Prism 8.

## Supporting information

Supplemental Materials List

Supplementary (raw) data 1

## Competing Interest Statement

The authors declare no competing interests.

## Acknowledgments

Some worm strains were kindly provided by the Caenorhabditis Genetics Center. We thank Dr. Lino Tessarollo for providing the *Tamalin* mutant mice, Justin Ruiz for technical support, Dr. Verena Jantsch for the α-pSer8 SUN-1 antibody, and Dr. Monique Zetka for the α-HIM-3 and α-HTP-3 antibodies, and members of the Colaiacovo laboratory for critical reading of this manuscript. This work was supported by National Institutes of Health grant R01GM072551 to M.P.C.

## Author Contributions

M.M.-G., P.R.N., P.W.J., and M.P.C conceived the study. M.M.-G., P.R.N., M.W.S., K.A.B., S.N., N.S., C.G.S.-G., T.T.S., S.B.-S., A.C., and S.P., performed the experiments. M.M.-G, P.R.N., M.W.S., and A.C., analyzed the data. M.M.-G., P.W.J., and M.P.C. wrote the original draft of the manuscript. M.M.-G., P.R.N., C.G.S.-G., T.T.S, A.C., E.M.-P., P.W.J. and M.P.C. reviewed and edited the manuscript. M.P.C. acquired the funding. P.W.J. and M.P.C. supervised the study. All authors read, reviewed, and approved the manuscript.

## Notes

### Competing Interest Statement

The authors have declared no competing interest.

