## Supplemental Materials List for "GRAS-1 is a conserved novel regulator of early meiotic chromosome dynamics in *C. elegans*"

**Supplemental Data. Raw data**

**Supplemental figure S1. Expression of MmCytip, MmGrasp, and Cegras-1 during spermatogenesis, and analysis of GRAS-1 localization and mutant phenotypes.**

**Supplemental figure S2. Switch in PLK-2 localization and SC central region formation are GRAS-1-dependent, but SC defects are IMA-1 and IMA-2-independent.**

**Supplemental figure S3. GRAS-1 interacts with SC central region proteins SYP-1, SYP-2, and SYP-3, and contributes to SPO-11-dependent DSB repair.**

**Supplemental figure S4. Tamalin-Cytip DKO male mouse analysis.**

**Supplemental figure S5. Tamalin-Cytip DKO female mouse analysis and HsCYTIP complementation of *gras-1* mutants.**

**Supplemental table 1. List of C. elegans lines used in this study**

**Supplemental table 2. Primary antibodies used for C. elegans immunostainings.**

**Supplemental table 3. Mouse genotyping primers and products**

**Supplemental table 4. Primary and secondary antibodies used for mouse chromatin spreads**

**Supplemental video 1. *gras-1* movement defects depend on dynein**

**Supplemental video 2. GRAS-1 phosphorylation affects its role in chromosome movement**
