## Supplemental Table 1 for "GRAS-1 is a conserved novel regulator of early meiotic chromosome dynamics in *C. elegans*"

**Supplemental table 1. List of *C. elegans* lines used in this study**

| Strain name | Genotype | Reference |
|---|---|---|
| **PHX598** | *gras-1::gfp(syb598[gras-1::gfp])(I)* | This study |
| **CV392** | *gras-1(rj15) I/hT2[bli-4(e937) let-? (q782) qls48] (I:III)* | This study |
| **CV464** | *gras-1(rj27) I/hT2[bli-4(e937) let-? (q782) qls48] (I:III)* | This study |
| **CV464** | *gras-1(rj28) I/hT2[bli-4(e937) let-? (q782) qls48] (I:III)* | This study |
| **CV826** | *gras-1(syb1380)/hT2[bli-4(e937) let-?(q782) qls48] (I;III)* | This study |
| **CV849** | *gras-1(syb1899)/hT2[bli-4(e937) let-?(q782) qls48] (I;III)* | This study |
| **AV106** | *spo-11 (ok79) IV/nT1 [unc-? (n754) let-?] (IV;V)* | (Dernburg et al. 1998) |
| **CV819** | *gras-1(rj28) I/hT2[bli-4(e937) let-? (q782) qls48] (I:III); spo-11 (ok79) IV/nT1 [unc-? (n754) let-?] (IV;V)* | This study |
| **CV835** | *oxls279[Ppie-1::GFP::H2B, unc +](II); ieSi21 [sun-1p::sun-1::mRuby::sun-1 3'UTR + Cbr-unc-119(+)] IV.* | This study; (Frøkjær-Jensen et |

| | | |
|---|---|---|
| | | al. 2008; Rog and Dernburg 2015) |
| **CV832** | *gras-1(rj28) I/hT2[bli-4(e937) let-? (q782) qls48] (I:III); oxls279[Ppie-1::GFP::H2B, unc +](II); ieSi21 [sun-1p::sun-1::mRuby::sun-1 3'UTR + Cbr-unc-119(+)] IV.* | This study |
| **CV812** | *gras-1::gfp(syb598[gras-1::gfp])(I); syp-2 (ok307) V/ nT1[Unc-? (n754) let-? qIs 50] (IV;V)* | This study; (Colaiácovo et al. 2003) |
| **CV818** | *gras-1::gfp(syb598[gras-1::gfp])(I); spo-11 (ok79) IV/nT1 [unc-? (n754) let-?] (IV;V)* | This study |
| **CV821** | *gras-1::gfp(syb598[gras-1::gfp])(I); chk-2(ok3037)* | This study; VC3236 |
| **WBM1119** | *wbmls60[pie-1p::3xFLAG::dpy-10 crRNA::unc-54 3'UTR, III]* | (Silva-García et al. 2019) |
| **CV882** | *gras-1(rj28) (I); wbmls60[pie-1p::3xFLAG::dpy-10 crRNA::unc-54 3'UTR, III]* | This study |
| **CV870** | *gras-1(rj28) (I); rj55[pie-1p::3XFLAG::HsCYTIP::unc-54 3'UTR, III]* | This study |
| **CA1199** | *ieSi38 [sun-1p::TIR1::mRuby::sun-1 3'UTR + Cbr-unc-119(+)] IV* | (Zhang et al. 2015) |
| **CA1215** | *dhc-1(ie28[dhc-1::degron::GFP]) I; ieSi38 [sun-1p::TIR1::mRuby::sun-1 3'UTR + Cbr-unc-119(+)] IV* | (Zhang et al., 2015) |
