## Supplemental Table 2 for "GRAS-1 is a conserved novel regulator of early meiotic chromosome dynamics in *C. elegans*"

**Supplemental table 2. Primary antibodies used for *C. elegans* immunostainings.**

| Antibody | Host | Dilution | Reference |
|---|---|---|---|
| α-GFP | Chicken | 1:500 | ab13970, Abcam |
| α-RFP | Rabbit | 1:100 | ab62341, Abcam |
| α-pSer8 SUN-1 | Guinea Pig | 1:700 | (Woglar et al. 2013) |
| α-HIM-8 | Rabbit | 1:500 | 41980002, Novus Biological - SDI |
| α-RAD-51 | Rabbit | 1:10,000 | 29480002, Novus Biological - SDI |
| α-HTP-3 | Guinea Pig | 1:500 | (Goodyer et al. 2008) |
| α-SYP-1 | Goat | 1:2,000 | (Nadarajan et al. 2017) |
| α-PLK-2 | Rabbit | 1:200 | (Nishi et al. 2008) |
| α-REC-8 | Rabbit | 1:500 | SDQ0802, Novus Biologicals |
| α-SYX-4 | Rabbit | 1:300 | (Jantsch-Plunger and Glotzer 1999) |
| Phalloidin-Atto 488 | - | 1:400 | 49409, Sigma Aldrich |
| α-HIM-3 | Chicken | 1:400 | Gift from M. Zetka (Goodyer et al. 2008) |
| α-ZHP-3 | Guinea Pig | 1:500 | (Bhalla et al. 2008) |
