## Supplemental Table 3 for "GRAS-1 is a conserved novel regulator of early meiotic chromosome dynamics in *C. elegans*"

**Supplementary table 3. Mouse genotyping primers and products**

| ***Gene*** | **Forward Primer (5'-…-3')** | **Reverse Primer (5'-...-3')** | **Band Size (bp)** |
|---|---|---|---|
| *Spo11-Cre* Transgene | CCATCTGCCA CCAGCCAG | TCGCCATCTTC CAGCAGG | 281 |
| *Cre* Internal Control (Cpxm1) | ACTGGGATCTTCG AACTCTTTGGAC | GATGTTGGGGCA CTGCTCATTCACC | 420 |
| *Tamalin* WT | CTGCTTGCAGGTTT CCACAGCTTC | CTACAGCCTTCTGA GACCCGAGTG | 330 |
| *Tamalin KO* | CTGCTTGCAGGTT TCCACAGCTTC | GAATGATGGCCTT AGTGGTTCGTG | 436 |
| *Cytip tm1b* | GCTACCATTACCAGTTGGT CTGGTGTC | TGAGTAGCTGGGA AGACCAATGTCC | 676 |
| *Cytip WT, Cytip tm1c* | CAATCCCACCAGA TCATCAACAGCC | ACTGATCATCCTGTCTCAGG TGTGG | 719, 861 |
| *Cytip tm1c* | GAGATGGCGCAACGCAATT AATG | TGAGTAGCTGGGAAGACCAA TGTCC | 335 |
