## Supplemental Table 4 for "GRAS-1 is a conserved novel regulator of early meiotic chromosome dynamics in *C. elegans*"

**Supplementary table 4. Primary and secondary antibodies used for mouse chromatin spreads**

| Antibody | Host | Source | Catalog Number | IF Dilution |
|---|---|---|---|---|
| ATR | Goat | Santa Cruz | sc-1887 | 1:50 |
| CDK2 | Mouse | Santa Cruz | sc-6248 | 1:50 |
| CYTIP | Rabbit | Novus | NBP1-88946 | 1:200 |
| RAD51 | Mouse | Invitrogen | MA5-14419 | 1:200 |
| RPA2 | Rabbit | Protein Tech Group | 10412-1-AP | 1:500 |
| SYCP1 | Rabbit | Thermo | PA1-16763 | 1:1000 |
| SYCP3 | Rabbit | Novus | NB300-231 | 1:1000 |
| SYCP3 | Goat | Novus Biologicals | AF3750 | 1:100 |
| SYCP3 | Mouse | Santa Cruz | sc-74568 | 1:50 |
| yH2AX | Mouse | Thermo | MA1-2022 | 1:1000 |
| Goat IgG (H+L) Alexa Fluor 488 | Donkey | Invitrogen | A-11055 | 1:500 |
| Goat IgG (H+L) Alexa Fluor 633 | Donkey | Invitrogen | A-21082 | 1:500 |
| Rabbit IgG (H+L) Alexa Fluor 568 | Donkey | Invitrogen | A-10042 | 1:500 |

| | | | | |
|---|---|---|---|---|
| Mouse IgG (H+L)<br>Alexa Fluor 488 | Donkey | Invitrogen | A-21202 | 1:500 |
| Mouse IgG (H+L)<br>Alexa Fluor 568 | Donkey | Invitrogen | A-11031 | 1:500 |
| Rabbit IgG (H+L)<br>Alexa Fluor 568 | Goat | Invitrogen | A-11011 | 1:500 |
| Mouse IgG (H+L),<br>Alexa Fluor 488 | Goat | Invitrogen | A-11001 | 1:500 |
