## Supplemental Figure S1 for "GRAS-1 is a conserved novel regulator of early meiotic chromosome dynamics in *C. elegans*"

Martinez-Garcia_FigS1

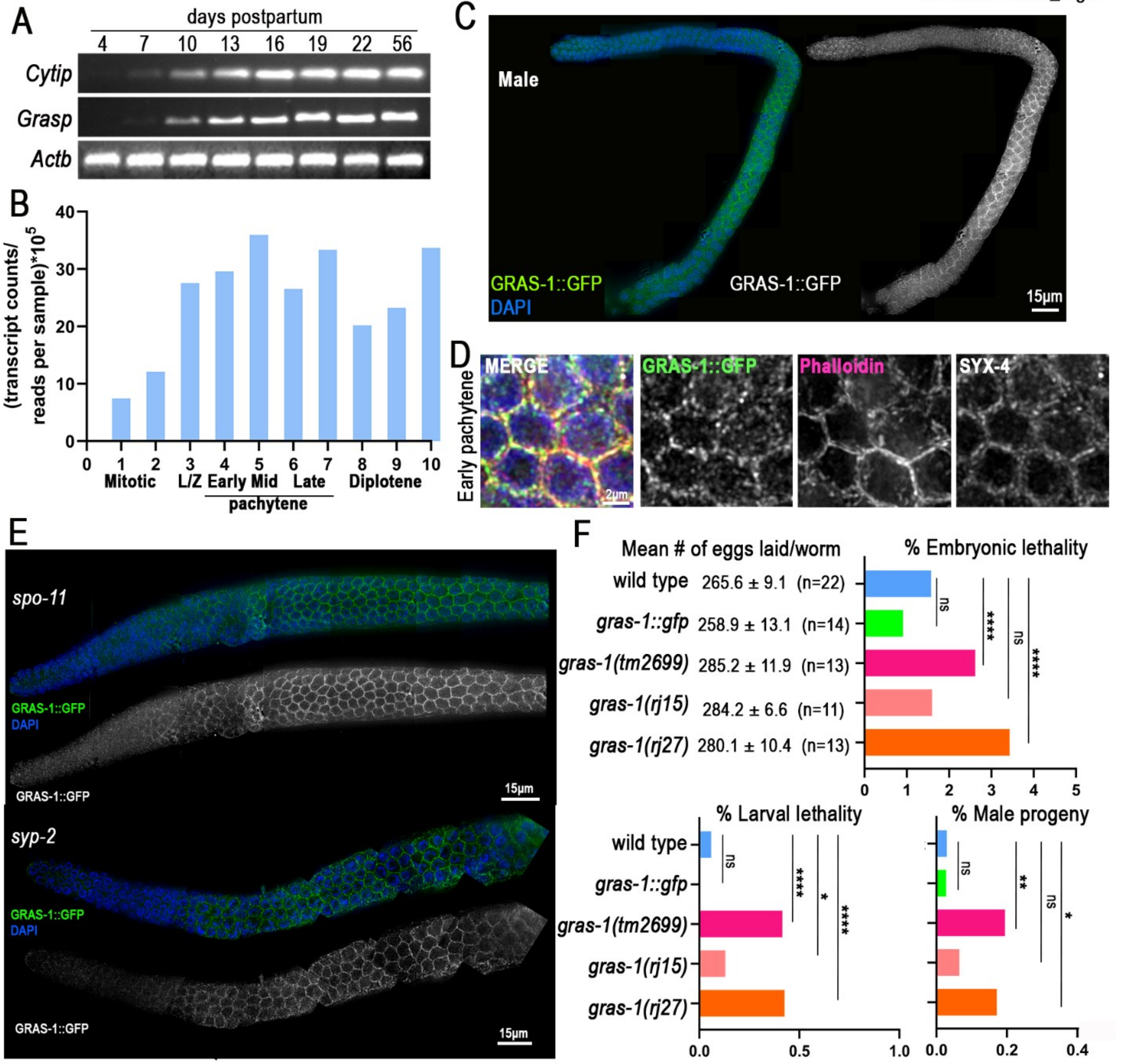

**Fig. S1. Expression of MmCytip, MmGrasp, and Ce*gras-1* during spermatogenesis, and analysis of GRAS-1 localization and mutant phenotypes. (A)** Expression of *Mus musculus* Cytip and Grasp during the first wave of spermatogenesis via RT-PCR. **(B)** Expression of *C. elegans gras-1* throughout the germline (zones 1-10) as described in (Tzur et al., 2017). **(C)** GRAS-1::GFP localization in whole mounted gonads of *gras-1::gfp* male *C. elegans* by co-staining with anti-GFP (green) and DAPI (blue). **(D)** Representative image of the early pachytene region in *gras-1::gfp* hermaphrodites co-stained for GRAS-1::GFP (green), Phalloidin (red), SYX-4 (white) and DAPI (blue). **(E)** GRAS-1::GFP localization in whole mounted gonads of *gras-1::gfp;spo-11* (top) *and gras-1::gfp;syp-2* (bottom) hermaphrodite *C. elegans* by co-immunostaining with anti-GFP (green) and DAPI (blue). GRAS-1::GFP signal alone is shown in white. **(F)** The mean number of eggs laid (brood size) ± SEM, as well as the percentage of embryonic lethality, larval lethality, and males are shown for the indicated genotypes. *$p < 0.05$, **$p < 0.01$, ****$p < 0.0001$ by Fisher's exact test. n= number of worms for which entire broods were analyzed in at least two independent biological replicates.
