## Supplemental Figure S2 for "GRAS-1 is a conserved novel regulator of early meiotic chromosome dynamics in *C. elegans*"

Martinez-Garcia_FigS2

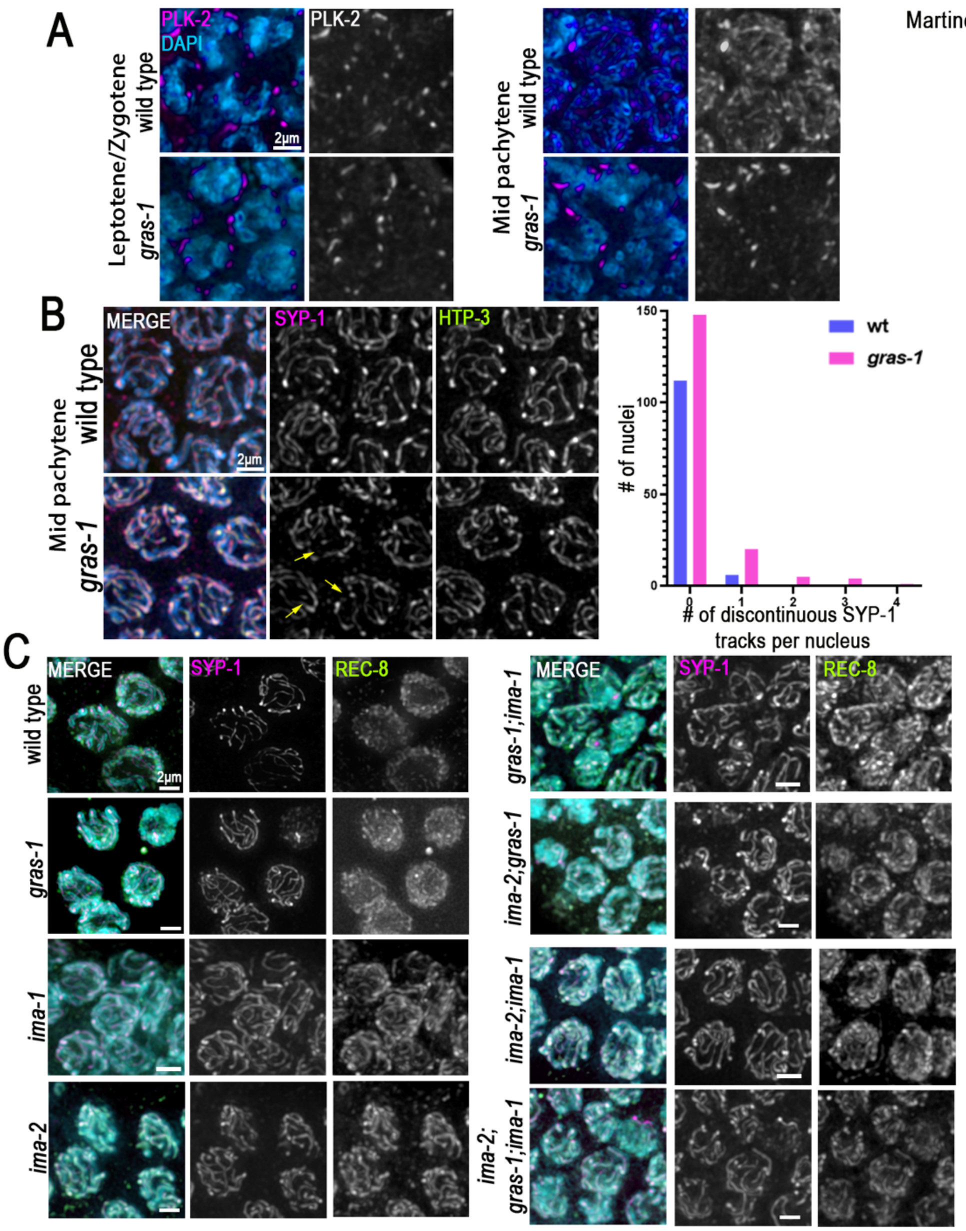

**Fig. S2. Switch in PLK-2 localization and SC central region formation are GRAS-1-dependent, but SC defects are IMA-1 and IMA-2-independent. (A)** High-resolution images of leptotene/zygotene and mid pachytene nuclei from wild type and *gras-1* germlines stained with anti-PLK-2 (magenta) and DAPI (blue). n=10 and 7 gonads each. **(B)** Left, high-resolution images of pachytene nuclei in wild type and *gras-1* germlines co-stained with anti-SYP-1 (magenta), anti-HTP-3 (green) and DAPI (blue). Yellow arrows indicate nuclei with SYP-1 discontinuities. Right, histogram showing the frequency of nuclei found with 0-4 SYP-1 discontinuities during mid pachytene stage in wild type and *gras-1*. n=118 and 178, respectively, p=0.002, Fisher's Exact test. **(C)** High-resolution images of mid pachytene stage nuclei from the indicated genotypes co-stained with anti-SYP-1 (magenta), anti-REC-8 (green) and DAPI (blue).
