## Supplemental Figure S3 for "GRAS-1 is a conserved novel regulator of early meiotic chromosome dynamics in *C. elegans*"

Martinez-Garcia_FigS3

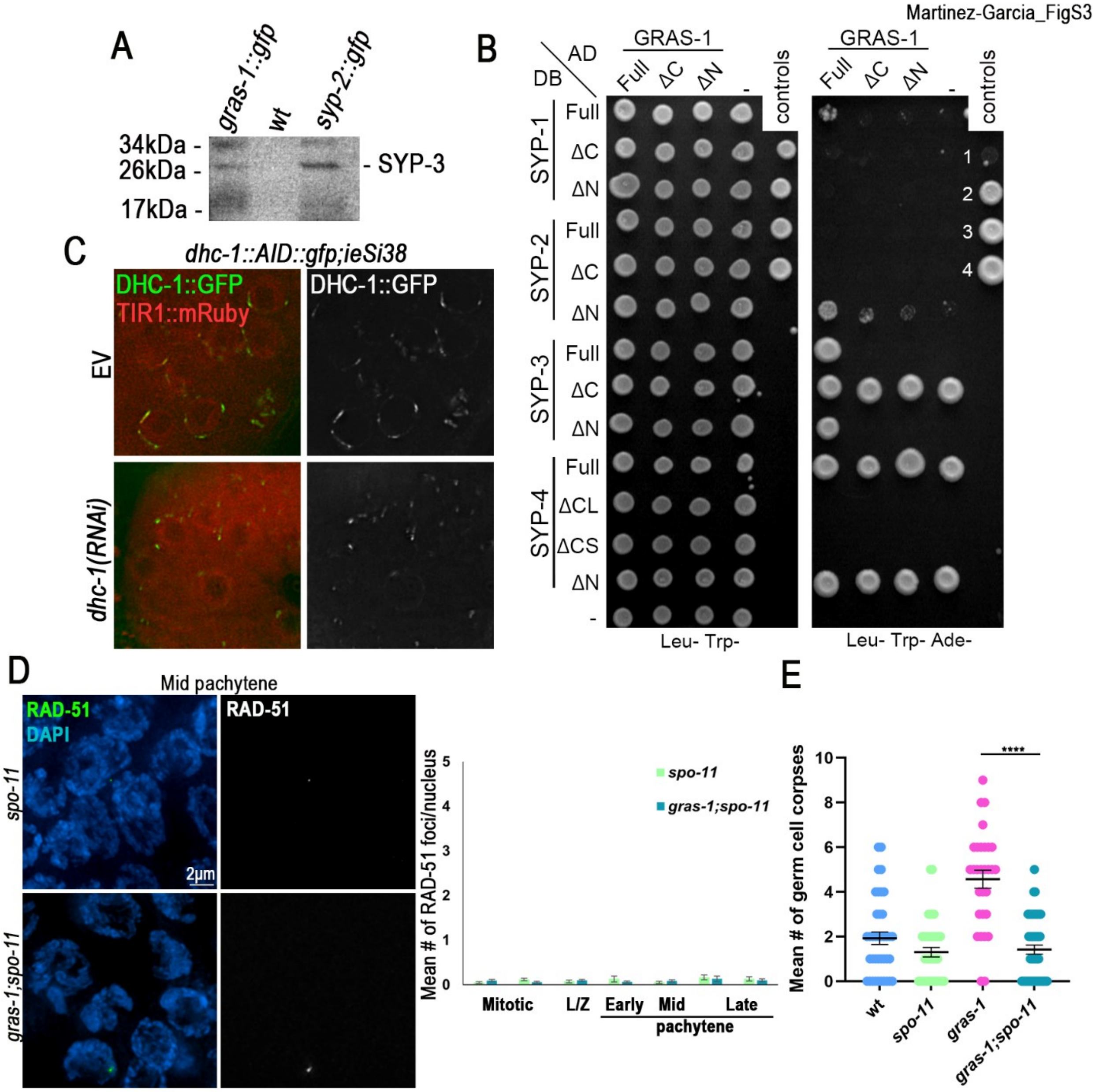

**Fig. S3. GRAS-1 interacts with SC central region proteins SYP-1, SYP-2, and SYP-3, and contributes to SPO-11-dependent DSB repair. (A)** Western blot using an anti-SYP-3 antibody showing immunoprecipitation of SYP-3 from *gras-1::gfp* and *syp-2::gfp* (positive control) but not wild type (negative control) whole worm lysates done with a GFP antibody. **(B)** The yeast two-hybrid system was used to examine the protein interactions between GRAS-1 full length, $\Delta Nt^{69-245}$ and $\Delta Ct^{1-163}$ truncations, and SYP-1/2/3/4 full length, N-terminal, and C-terminal truncations (AD, activation domain; DB, Gal4 DNA binding domain). Negative (no. 1) and positive controls (nos. 2-4) were used as described in (Schild-Prüfert et al. 2011). SYP-3 ΔC, SYP-4 full length and SYP-4 ΔN exhibited strong self-activation and therefore their observed interactions are false positives. **(C)** Live imaging of DHC-1::GFP (green) and TIR1::mRuby (red) proteins in leptotene/zygotene nuclei *of dhc-1::AID::gfp;ieSi38* worms grown in bacteria expressing either the empty vector or *dhc-1(RNAi).* **(D)** Left, high-resolution images of mid-pachytene nuclei in *spo-11* and *gras-1;spo-11* stained with anti-RAD-51 (green) and DAPI (blue). Right, Histogram showing the mean number of RAD-51 foci/nucleus scored along the germlines of the indicated genotypes. 6 gonads were scored per genotype in two independent biological replicates. Error bars represent the SEM. Not significant by the Mann-Whitney U-test. **(E)** Histogram showing the mean number of germ cell corpses in wild type, *spo-11*, *gras-1,* and *gras-1;spo-11* worms. Error bars represent the SEM. ****$p<0.0001$, Mann-Whitney U-test, n= 39, 40, 30 and 48 gonads, respectively.
