## Supplemental Figure S4 for "GRAS-1 is a conserved novel regulator of early meiotic chromosome dynamics in *C. elegans*"

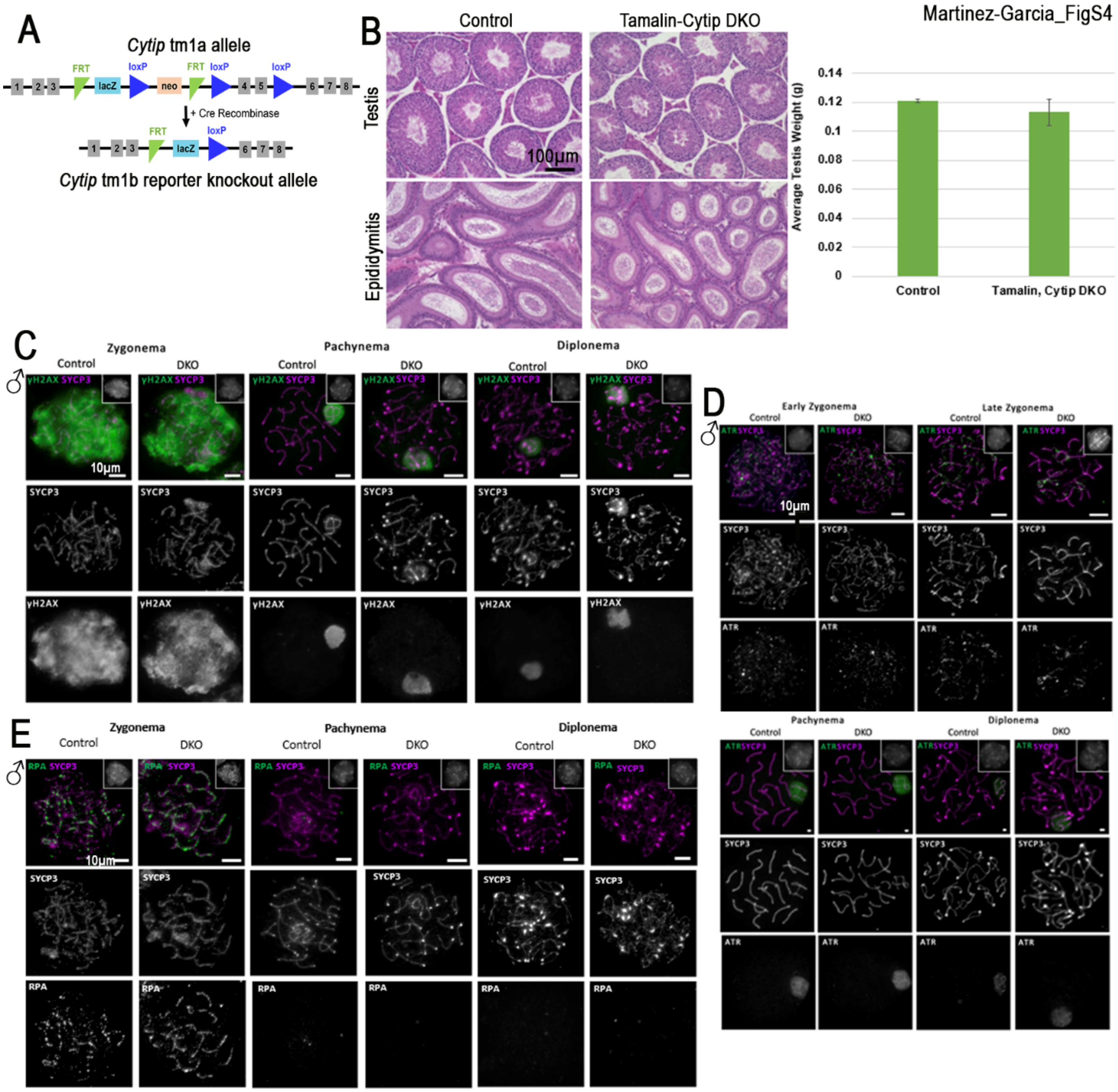
Martinez-Garcia_FigS4
A
Cytip tm1a allele
FRT
loxP
lacZ
neo
+ Cre Recombinase
Cytip tm1b reporter knockout allele
B
Control
Tamalin-Cytip DKO
Testis
Epididymitis
100µm
Average Testis Weight (g)
0.14
0.12
0.1
0.08
0.06
0.04
0.02
0
Control
Tamalin, Cytip DKO
C
Zygonema
Pachynema
Diplonema
Control
DKO
γH2AX SYCP3
SYCP3
γH2AX
10µm
D
Early Zygonema
Late Zygonema
ATR SYCP3
ATR
E
RPA SYCP3
RPA

**Fig. S4. Tamalin-Cytip DKO male mouse analysis. (A)** Representative structure of *Cytip* tm1a and tm1b *Mus musculus* alleles. Numbered exons shown in grey boxes, FRT: flippase recognition target, lacZ reporter, loxP: locus of X-over P1 site, neo: neomycin resistance gene. **(B)** Cross section of testis (top panel) and epididymitis (bottom panel) of control (301dpp) and Tamalin-Cytip DKO (196dpp) mice stained with hematoxylin and eosin. Graph shows no significant difference in testis weight from control and Tamalin-Cytip DKO. Error bars show mean ± SEM. Two-tailed Student's t-test, n=3 mice each**. (C)** Chromatin spreads from early meiotic prophase (zygonema), mid meiotic prophase (pachynema) and late meiotic prophase (diplonema) control and *Tamalin-Cytip* DKO *Mus musculus* spermatocytes co-immunostained with antibodies against SYCP3 (magenta) and γ-H2AX (green). Insets show normal chromatin morphology (DAPI). n=50 cells per mouse and 3 mice per genotype. **(D)** Chromatin spreads from early and late zygonema, pachynema and diplonema cells of control and *Tamalin-Cytip* DKO *Mus musculus* spermatocytes co-immunostained with antibodies against SYCP3 (magenta) and ATR (green). Insets show normal chromatin morphology (DAPI). **(E)** Chromatin spreads from early and late zygonema, pachynema and diplonema cells of control and *Tamalin-Cytip* DKO *Mus musculus* spermatocytes co-immunostained with antibodies against SYCP3 (magenta) and RPA (green). Insets show normal chromatin morphology (DAPI).
