## Supplemental Figure S5 for "GRAS-1 is a conserved novel regulator of early meiotic chromosome dynamics in *C. elegans*"

Martinez-Garcia_FigS5

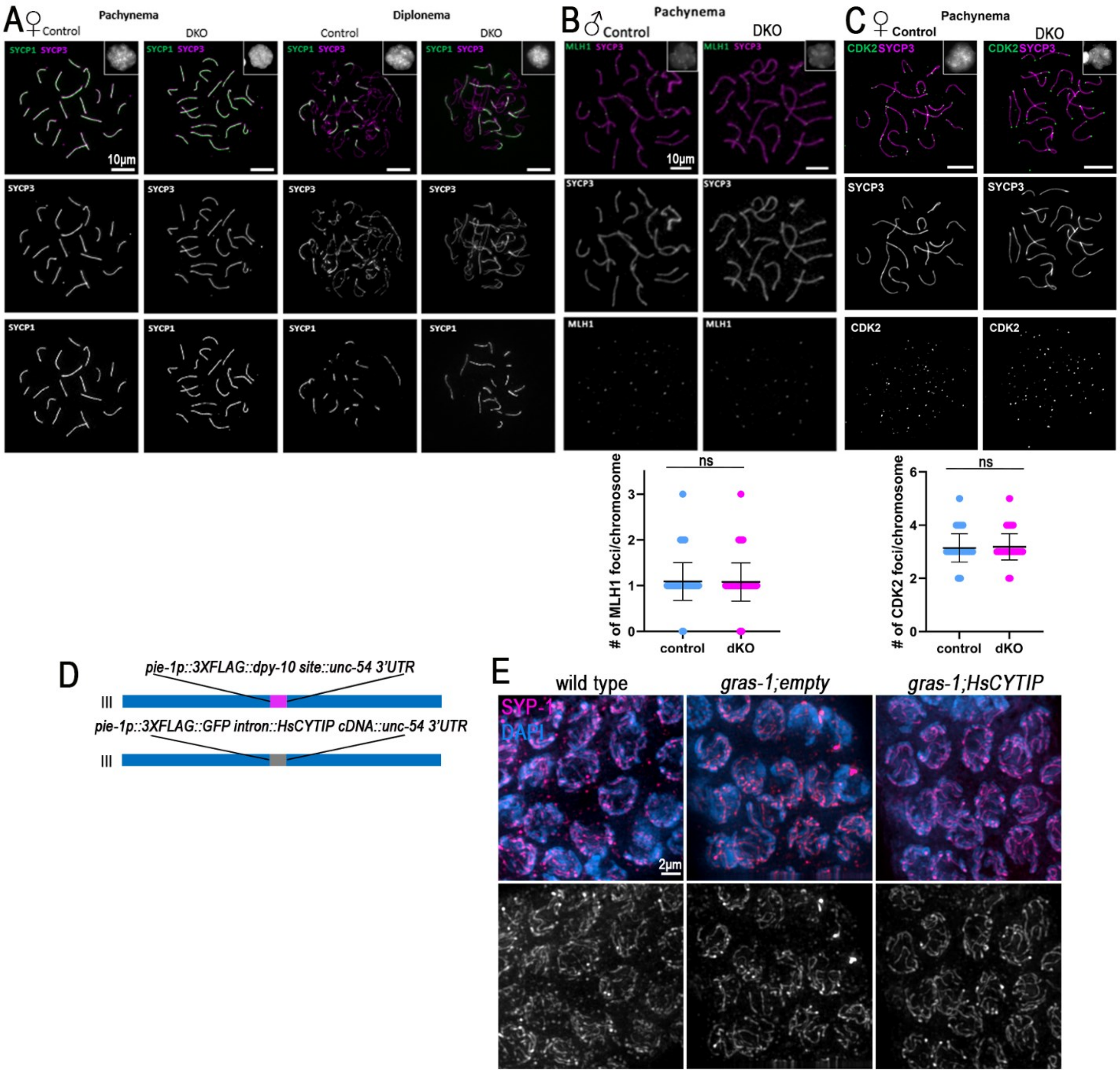

**Fig. S5. Tamalin-Cytip DKO female mouse analysis and HsCYTIP complementation of *gras-1* mutants. (A)** Chromatin spreads from pachynema and diplonema cells of control and *Tamalin-Cytip* DKO *Mus musculus* oocytes co-immunostained with antibodies against SYCP1 (green) and SYCP3 (magenta). Insets show normal chromatin morphology (DAPI). **(B)** Top, chromatin spreads from pachynema cells of control and *Tamalin-Cytip* DKO *Mus musculus* spermatocytes co-immunostained with antibodies against SYCP3 (magenta) and MLH1 (green). Insets show normal chromatin morphology (DAPI). Bottom, dot plot of the number of MLH1 foci per chromosome quantified in control and DKO oocytes. 21 cells per genotype, ns: not significant by Mann-Whitney U-test. **(C)** Top, chromatin spreads from pachynema cells of control and *Tamalin-Cytip* DKO *Mus musculus* oocytes co-immunostained with antibodies against SYCP3 (magenta) and CDK2 (green). Insets show normal chromatin morphology (DAPI). Bottom, dot plot of the number of CDK2 foci per chromosome quantified in control and DKO oocytes. 25 cells per genotype, ns: not significant by Mann-Whitney U-test. **(D)** Schematic representation of the genomic location of SKI LODGE germline insertion and the HsCYTIP complementation cassette. **(E)** High-resolution images of whole mounted gonads of wild type, *gras-1;empty* and *gras-1;HsCYTIP* during early pachytene co-stained with anti-SYP-1 (magenta) and DAPI (blue).
